# A new class of polymorphic T6SS effectors and tethers

**DOI:** 10.1101/2022.10.27.514009

**Authors:** Katarzyna Kanarek, Chaya Mushka Fridman, Eran Bosis, Dor Salomon

**Author notes:** Address correspondence to Dor Salomon, and to Eran Bosis.

## Abstract

Bacteria use the type VI secretion system (T6SS) to deliver toxic effectors into bacterial or eukaryotic cells during interbacterial competition, host colonization, or when resisting predation. The identity of many effectors remains unknown. Here, we identify RIX, a new domain that defines a class of polymorphic T6SS cargo effectors. RIX, which is widespread in the *Vibrionaceae* family, is located at N-termini of proteins containing diverse antibacterial and anti-eukaryotic toxin domains. We demonstrate that RIX-containing proteins are delivered via T6SS into neighboring cells, and that RIX is necessary and sufficient for secretion. We show that RIX-containing proteins can also act as tethers, enabling the T6SS-mediated delivery of other cargo effectors by a previously undescribed mechanism. RIX-containing proteins significantly enlarge the repertoire of known T6SS effectors, especially those with anti-eukaryotic activities. Our findings also suggest that T6SSs may play a major, currently underappreciated, role in interactions between vibrios and eukaryotes.

## Introduction

The type VI secretion system (T6SS) is a protein delivery apparatus in Gram-negative bacteria, which was originally described as an anti-eukaryotic determinant ^1,2^. Nevertheless, further investigations revealed that most T6SSs play a role in interbacterial competition ^3,4^, whereas only a few T6SSs have been identified as anti-eukaryotic ^5,6^. These roles are mediated by toxic proteins, called effectors, which are deployed inside neighboring bacterial and eukaryotic cells_^5,7,8^_.

Effectors are loaded onto a missile-like structure, which is ejected by a contractile sheath that engulfs it in the cytoplasm of the secreting cell ^9,10^. The missile is composed of Hcp proteins that are stacked as hexameric rings forming an inner tube; the tube is capped by a spike complex comprising a VgrG trimer sharpened by a PAAR repeat-containing protein (hereafter referred to as PAAR) ^9^. T6SS effectors can be divided into two types: (1) specialized effectors, which are secreted structural components (i.e., Hcp, VgrG, or PAAR) containing a C-terminal toxin domain extension ^11–14^ and (2) cargo effectors, which are toxic proteins that non-covalently interact with one of the missile components or its C-terminal extension ^15–19^, either directly or aided by an adaptor protein ^20,21^ or a co-effector ^22^. Because cargo effectors can bind to diverse loading platforms on the tube or spike, they lack a canonical secretion signal or domain. Therefore, identifying cargo effectors is a challenging task, especially if they are not encoded within T6SS gene clusters or near T6SS-associated genes.

Many T6SS cargo effectors belong to one of three known classes of polymorphic toxins: (1) MIX (Marker for type sIX) domain-containing proteins ^15,23^; (2) FIX (Found in type sIX) domain-containing proteins ^16^; or (3) Rhs (Rearrangement hotspot) repeat-containing proteins ^24–28^. These three domains are found N-terminal to diverse C-terminal toxin domains, and they are predicted to play a role in their delivery. MIX and FIX domains are specifically found in T6SS-secreted proteins, whereas Rhs repeats are also present in toxins secreted by other types of secretion systems. However, many other cargo effectors lack a known domain at their N-terminus; it is possible that these effectors contain N-terminal delivery domains that have not yet been revealed.

We previously identified Tme1, an antibacterial T6SS effector of *Vibrio parahaemolyticus*. Tme1 contains a C-terminal toxin domain named Tme (T6SS Membrane-disrupting Effector), which permeabilizes membranes and dissipates membrane potential ^29^; the sequence N-terminal to the Tme domain does not contain a known domain or activity. Here, we show that the N-terminus of Tme1 is necessary and sufficient to mediate T6SS secretion in *V. parahaemolyticus*, and we use it to reveal a new domain that is widespread in members of the *Vibrionaceae* family. Proteins containing the identified domain, named RIX (aRginine-rich type sIX), are secreted via T6SS. RIX is found N-terminal to diverse C-terminal extensions with antibacterial and anti-eukaryotic toxic activities, as well as to sequences that function as loading platforms for cargo effectors. Therefore, we reveal a new class of T6SS-secreted proteins, including polymorphic toxins and effector tethers.

## Results

### The N-terminus of Tme1 is necessary and sufficient for T6SS-mediated secretion

We previously reported that Tme1 is an antibacterial effector delivered by T6SS1 in *V. parahaemolyticus* BB22OP ^29^; its toxin domain, Tme, is located at the C-terminus. We hypothesized that the N-terminus of Tme1 plays a role in secretion via T6SS. To test this hypothesis, we monitored the expression and secretion of a truncated version of Tme1 lacking the first 60 amino acids (Tme1^61-310^) (**Fig. 1A**). As shown in **Fig. 1B**, truncating the 60 N-terminal amino acids abrogated Tme1 secretion, indicating that this region is necessary for T6SS-mediated secretion. Next, we sought to determine whether the N-terminus of Tme1 is sufficient for T6SS-mediated secretion. To this end, we used Tse1, a *Pseudomonas aeruginosa* T6SS effector. Tse1 is unable to secrete via *V. parahaemolyticus* T6SS1 (**Fig. 1C**, right lanes). However, fusing the N-terminal 111 amino acids of Tme1 to Tse1 (Tme1^1-111^-Tse1) enabled its secretion via *V. parahaemolyticus* T6SS1 (**Fig. 1C**, left lanes). Shorter N-terminal Tme1 sequences were unstable when fused to Tse1, and were therefore not tested. Taken together, these results suggest that a region found at the N-terminus of Tme1 is necessary and sufficient for T6SS-mediated secretion.

**Fig. 1.**
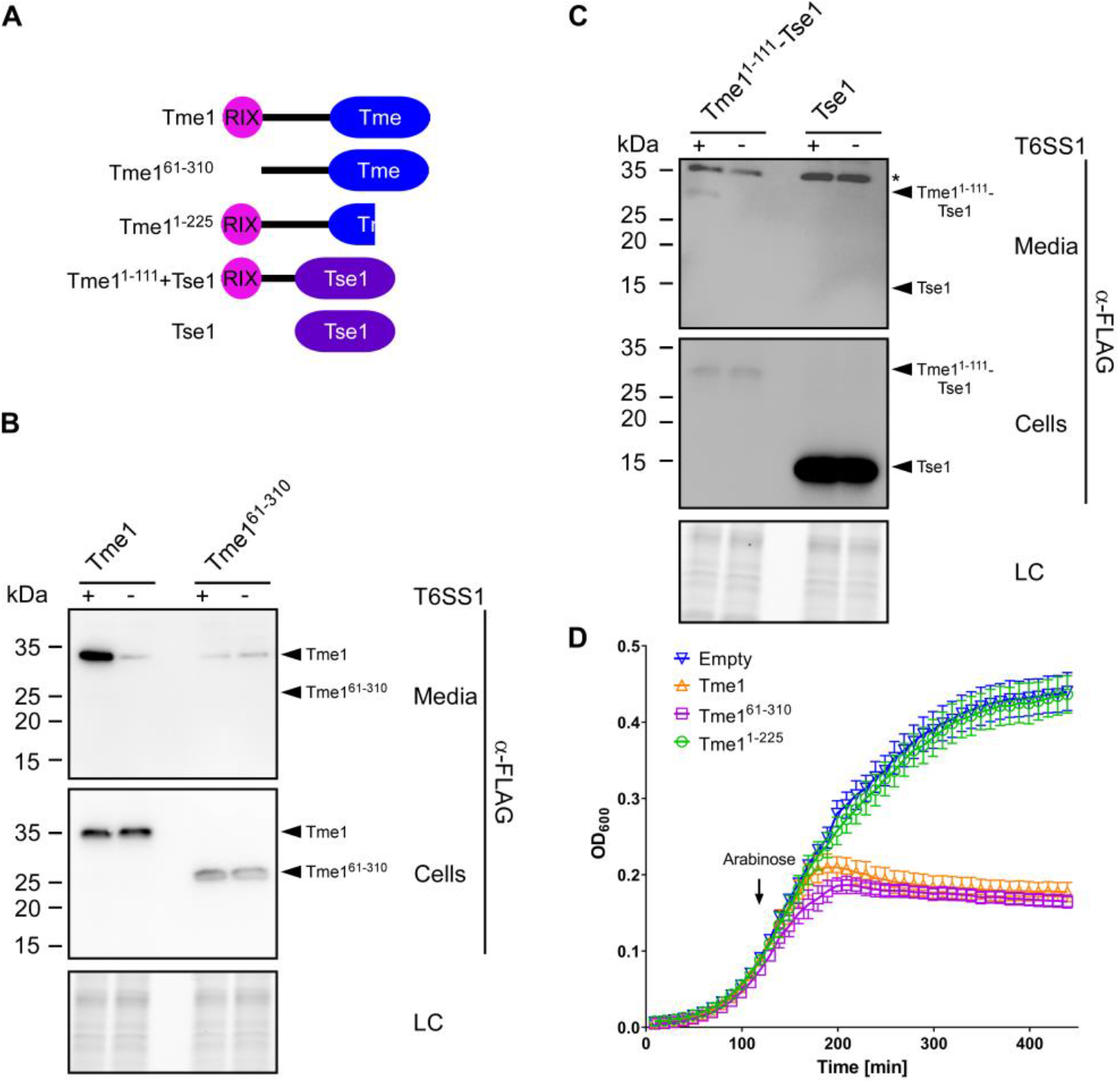
The N-terminus of Tme1 is necessary and sufficient for T6SS-mediated secretion in *V. parahaemolyticus*. **(A)** Schematic representation of Tme1 truncations and fusion proteins used in this figure. **(B-C)** Expression (cells) and secretion (media) of the indicated C-terminal FLAG-tagged proteins expressed from pBAD33.1^F^-based plasmids in *V. parahaemolyticus* BB22OP ∆*hns*/∆*tme1* (T6SS1^+^; deletion of *hns* was used to hyper-activate T6SS1) and *V. parahaemolyticus* BB22OP ∆*hns*/∆*tme1*/∆*hcp1* (T6SS1^−^). Samples were grown in MLB supplemented with chloramphenicol and 0.01% (wt/vol) L-arabinose (to induce expression from plasmids) for 3 h at 30°C. Loading control (LC) is shown for the total protein lysates. In (C), the asterisk denotes a non-specific band detected by the α-FLAG antibody. **(D)** Growth of *E. coli* BL21 (DE3) containing pPER5 plasmids for the arabinose-inducible expression of the indicated proteins fused to an N-terminal PelB signal peptide (for delivery to the periplasm). An arrow denotes the timepoint (120 min) at which arabinose (0.05% wt/vol) was added to the media.

To confirm that the N-terminus of Tme1 does not play a role in the effector’s antibacterial activity, we monitored the toxicity of Tme1 variants with truncations at the N-terminus and the C-terminus. As shown in **Fig. 1D**, truncating the first 60 amino acids of Tme1 did not affect the toxicity of this effector in the periplasm of *E. coli*, whereas truncating the last 85 amino acids of Tme1, corresponding to the end of the Tme domain, abrogated its toxicity. The expression of all Tme1 forms was detected in immunoblots (**Supplementary Fig. S1**). Thus, the N-terminal end of Tme1 is not required for toxicity.

### RIX is an arginine-rich domain found at N-termini of polymorphic toxins

Following these findings, we set out to identify regions homologous to the N-terminus of Tme1 in other proteins. We identified 502 unique protein accession numbers that contain a homologous sequence at their N-terminus (**Supplementary Dataset S1**). A multiple sequence alignment of these homologs revealed a conserved, arginine-rich motif corresponding to amino acids 1-55 in Tme1 (**Fig. 2A**); hereafter, we will refer to this region as the RIX (aRginine-rich type sIX) domain. RIX-containing proteins are encoded by 692 Gamma-proteobacterial strains, exclusively belonging to the marine bacteria families *Vibrionaceae* (i.e., *Vibrio* and *Photobacterium*), and *Moritellaceae* (i.e., *Moritella*) (**Fig. 2B** and **Supplementary Dataset S1**). Many of the bacterial strains encoding RIX-containing proteins are pathogens of humans and animals; these include *V. parahaemolyticus, V. cholerae, V. vulnificus, V. campbellii, V. coralliilyticus*, and *V. crassostreae* ^30–33^. Importantly, although none of the identified RIX-containing proteins is encoded within a T6SS gene cluster or module, almost all (97.7%) of the genomes encoding RIX-containing proteins harbor a T6SS (**Supplementary Dataset S2**).

**Fig. 2.**
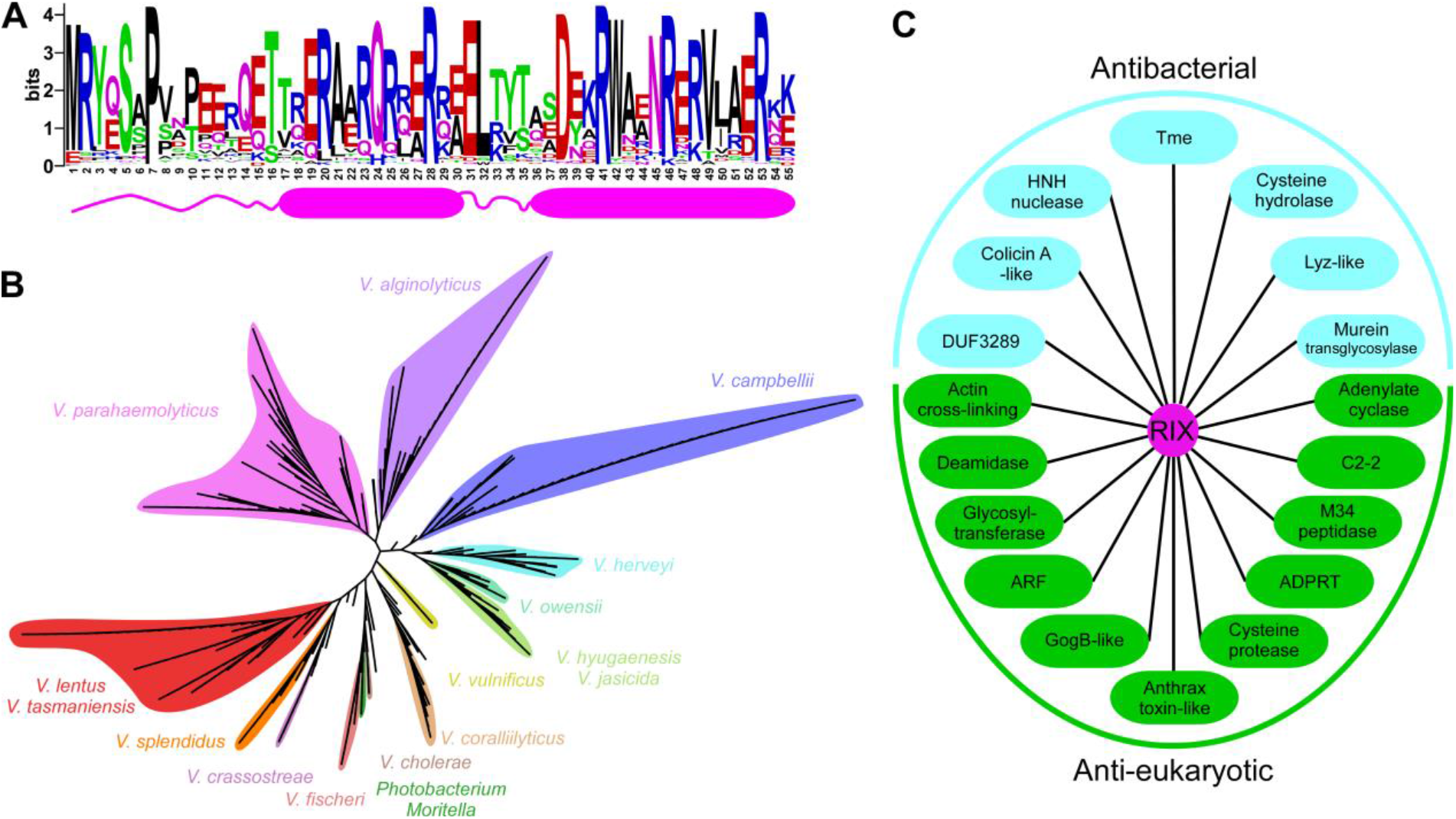
RIX is found in polymorphic toxins that are widespread in vibrios. **(A)** A conserved motif found at the N-terminus of polymorphic toxins (RIX) is illustrated using WebLogo, based on multiple sequence alignment of sequences homologous to the N-terminal 55 amino acids of Tme1. The position numbers correspond to the amino acids in Tme1. A secondary structure prediction, based on the AlphaFold2 prediction of the Tme1 structure, is shown below. Alpha helices are denoted by cylinders. **(B)** Phylogenetic distribution of bacteria encoding a protein with a RIX domain, based on the DNA sequence of *rpoB* coding for DNA-directed RNA polymerase subunit beta. The evolutionary history was inferred using the neighbor-joining method. **(C)** Examples of known and predicted activities and domains found in sequences C-terminal to RIX domains.

Analysis of the amino acid sequences C-terminal to RIX based on all-against-all pairwise similarity (using the CLANS classification tool ^34^) revealed 33 distinct clusters (**Table 1** and **Supplementary Fig. S2**). Remarkably, the majority of these C-terminal sequences contain domains that are known or predicted to be toxins with anti-eukaryotic activities (e.g., actin cross-linking, deamidase, adenylate cyclase, and glycosyltransferase) or antibacterial activities (e.g., pore-forming, HNH nuclease, and lysozyme-like) (**Fig. 2C** and **Table 1**). RIX-containing proteins with predicted antibacterial toxin domains are encoded upstream of a gene that possibly encodes a cognate immunity protein; genes encoding predicted anti-eukaryotic toxins do not neighbor a potential immunity gene. Notably, AlphaFold2 structure predictions ^35,36^ of representatives from each cluster revealed a possible conserved RIX structure, comprising two alpha helices preceded and connected by short loops (**Fig. 2A** and **Supplementary Fig. S3**).

**Table 1.**
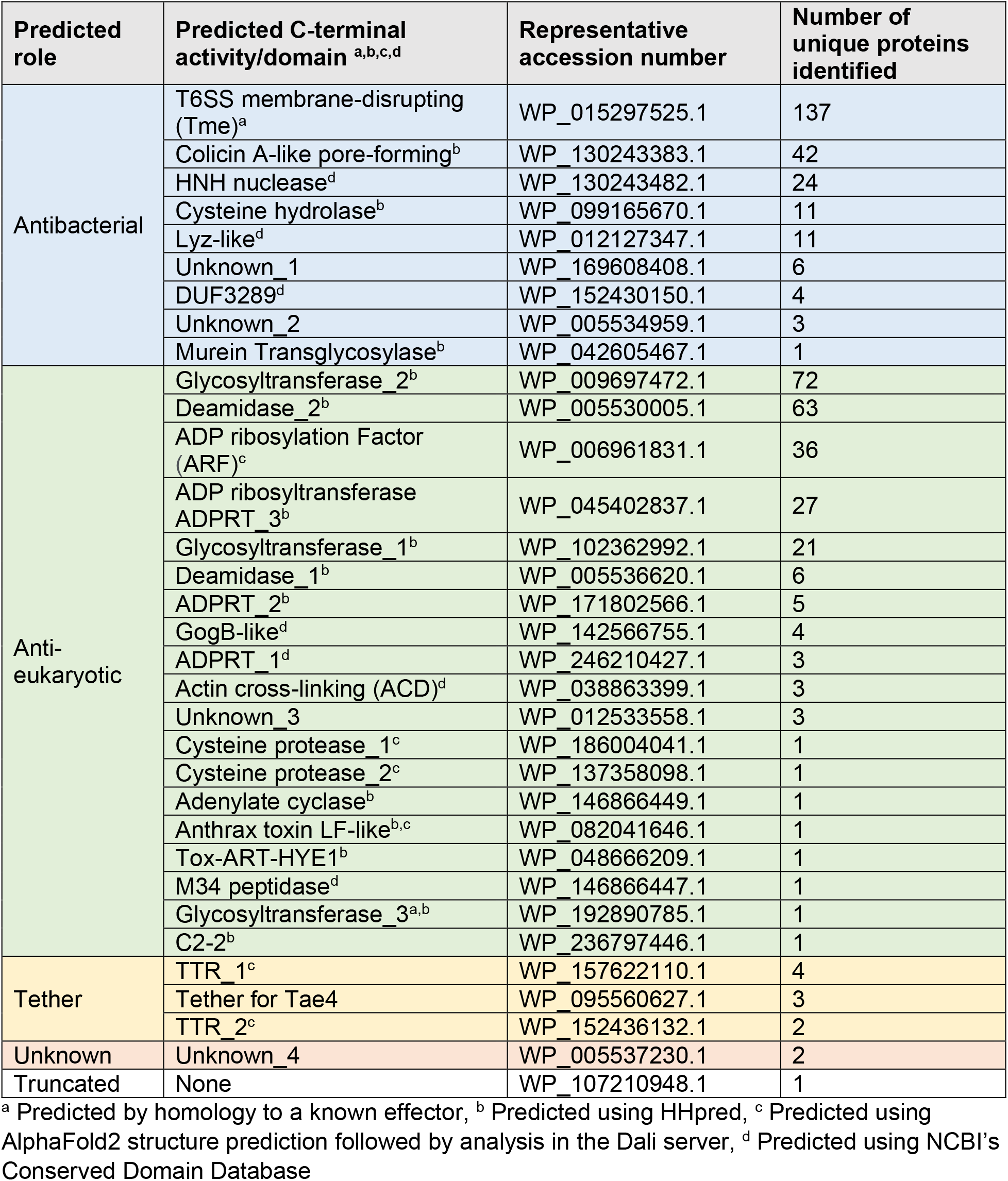
RIX-associated C-terminal extensions.

### RIX domain-containing proteins are secreted by T6SS

The results thus far indicated that: (1) Tme1 is a RIX-containing T6SS effector, (2) nearly all RIX-containing proteins are encoded by bacteria with a T6SS, and (3) most RIX-containing proteins have predicted C-terminal toxin domains. All together, these results led us to hypothesize that RIX-containing proteins are a new class of polymorphic T6SS effectors. To test this hypothesis, we investigated whether a predicted RIX-containing protein from *V. campbellii* ATCC 25920, WP_005534959.1, which has a C-terminal domain of unknown function (Unknown_2; **Table 1** and **Supplementary Fig. S4A**) functions as a T6SS effector. We first monitored the toxicity of this protein in bacteria. Expression of the WP_005534959.1 alone in *E. coli* was detrimental, whereas co-expression of the downstream-encoded protein (WP_005534960.1) antagonized this toxicity (**Supplementary Fig. S4B**). We then tested the T6SS-mediated delivery of the predicted effector using a previously established T6SS surrogate system in a *V. parahaemolyticus* RIMD 2210633 derivative strain ^29,37^. We found that expression of the predicted effector and immunity proteins from a plasmid in the surrogate attacker strain led to the killing of a parental prey strain lacking the predicted immunity protein (**Fig. 3A**). The killing was T6SS-dependent, since expression of the proteins in a derivative surrogate strain in which T6SS1 is inactive (∆*hcp1*) did not lead to killing of the prey strain. Expression of the predicted immunity protein from a plasmid in the prey strain protected it from this attack. These results indicate that the RIX domain-containing WP_005534959.1 and its downstream encoded WP_005534960.1 are an antibacterial T6SS effector and immunity pair.

**Fig. 3.**
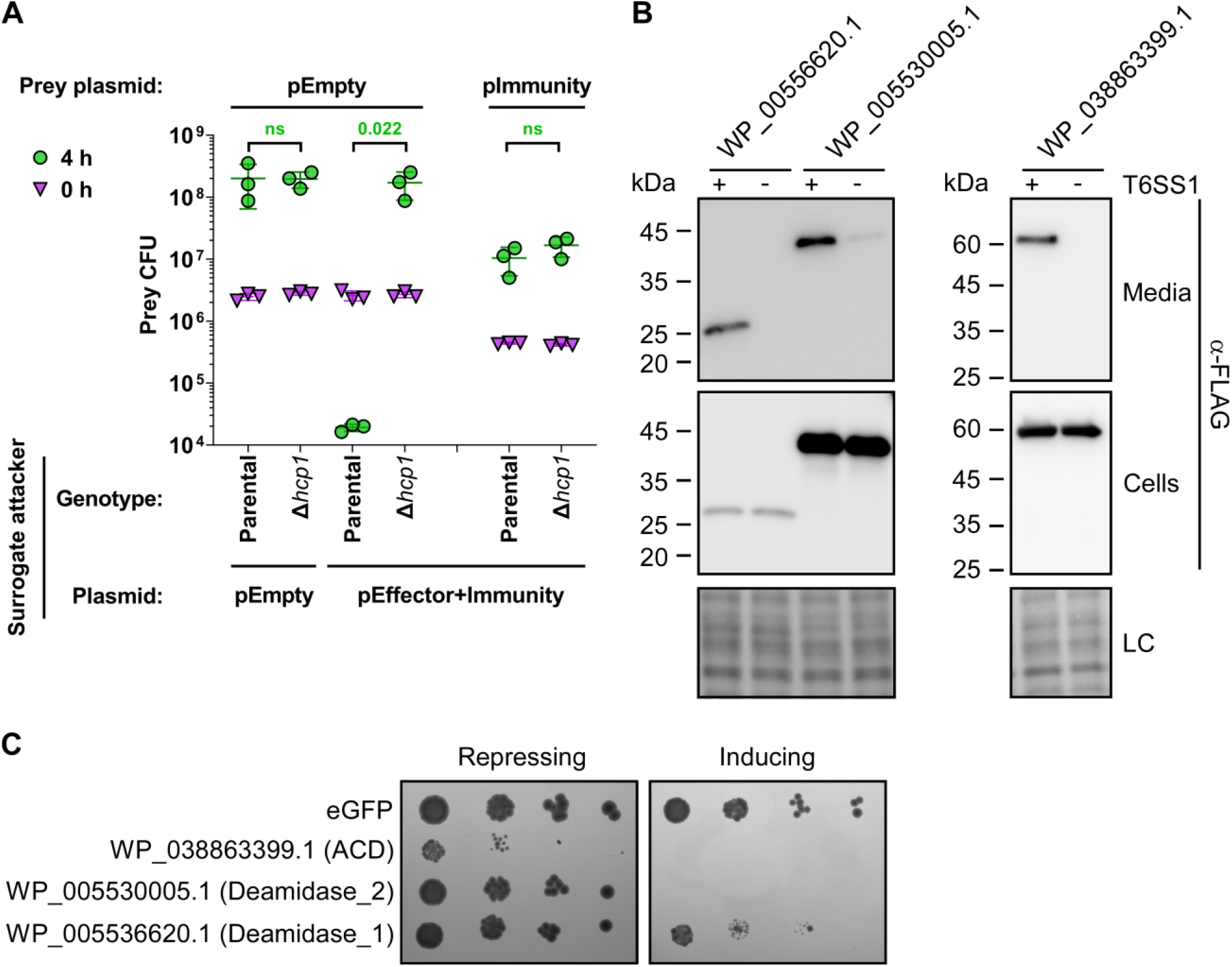
RIX-containing proteins are delivered and secreted by T6SS. **(A)** Viability counts (CFU) of *V. parahaemolyticus* RIMD 2210633 ∆*hcp1* prey strains harboring either an empty plasmid (pEmpty) or a plasmid for the arabinose-inducible expression of WP_005534960.1 (pImmunity) before (0 h) and after (4 h) co-incubation with a surrogate attacker strain (*V. parahaemolyticus* RIMD 2210633 derivative) or its T6SS1^−^ derivative (∆*hcp1*) carrying an empty plasmid or a plasmid for the arabinose-inducible expression of WP_005534959.1 and WP_005534960.1 (pEffector+Immunity). The statistical significance between samples at the 4 h timepoint was calculated using an unpaired, two-tailed Student’s *t*-test; ns, no significant difference (p>0.05). Data are shown as the mean ± SD; *n* = 3. **(B)** Expression (cells) and secretion (media) of the indicated C-terminal FLAG-tagged proteins expressed from pBAD33.1^F^-based plasmids in a *V. parahaemolyticus* RIMD 2210633-derivative surrogate (T6SS1^+^) strain or its ∆*hcp1* derivative (T6SS1^−^). Samples were grown in MLB supplemented with chloramphenicol and 0.05% (wt/vol) L-arabinose (to induce expression from plasmids) for 4 h at 30°C. Loading control (LC) is shown for the total protein lysates. **(C)** Tenfold serial dilutions of yeast strains carrying pGML10 plasmids for the galactose-inducible expression of the indicated C-terminal Myc-tagged protein were spotted on repressing (4% glucose) or inducing (2% wt/vol galactose and 1% wt/vol raffinose) agar plates.

To further support our hypothesis, we next determined whether RIX-containing proteins belonging to diverse clusters are secreted by T6SS. To this end, we monitored the expression and secretion of three additional RIX-containing proteins encoded by *V. campbellii* ATCC 25920: WP_005536620.1, WP_005530005.1, and WP_038863399.1; these proteins are predicted to have anti-eukaryotic toxin domains at their C-terminus (Deamidase_1, Deamidase_2, and Actin cross-linking, respectively; see **Table 1**). Notably, the genomic neighborhood of their encoding genes did not include T6SS-associated genes that would suggest that these RIX-containing proteins are T6SS effectors (**Supplementary Fig. S5** and **Dataset S1**). Remarkably, the three RIX-containing proteins were secreted from a surrogate strain in a T6SS-dependent manner (**Fig. 3B**). Furthermore, they were all toxic when expressed in the eukaryotic yeast model organism, *Saccharomyces cerevisiae* (**Fig. 3C**), suggesting that they affect a conserved eukaryotic target ^38^. Taken together, these results support our hypothesis that RIX-containing proteins are secreted by T6SSs.

Next, we sought to demonstrate the secretion of another RIX-containing protein via its endogenous T6SS, in addition to Tme1, which was shown previously (**Fig. 1B**). To this end, we investigated WP_157622110.1, encoded by *V. coralliillyticus* BAA-450. To monitor the secretion of WP_157622110.1 via T6SS in *V. coralliillyticus*, we set out to identify the conditions under which T6SS1 of *V. coralliilyticus* BAA-450 is active. First, we monitored the secretion of the T6SS spike protein VgrG1, and the ability of this strain to intoxicate *V. natriegens* prey bacteria during competition under different temperatures in media containing 3% (w/v) NaCl. Our results revealed that *V. coralliilyticus* BAA-450 T6SS1 is an antibacterial system that is active at 30°C (i.e., under warm, marine-like conditions) (**Supplementary Fig. S6**). We then monitored the secretion of the RIX-containing protein, WP_157622110.1, when expressed from a plasmid. As shown in **Fig. 4A**, WP_157622110.1 was secreted by *V. coralliilyticus* BAA-450 in a T6SS1-dependent manner, confirming the secretion of RIX-containing proteins via their endogenous T6SS. In addition, we confirmed that this RIX-containing protein is secreted in a T6SS-dependent manner from a surrogate strain (**Fig. 4B**).

**Fig. 4.**
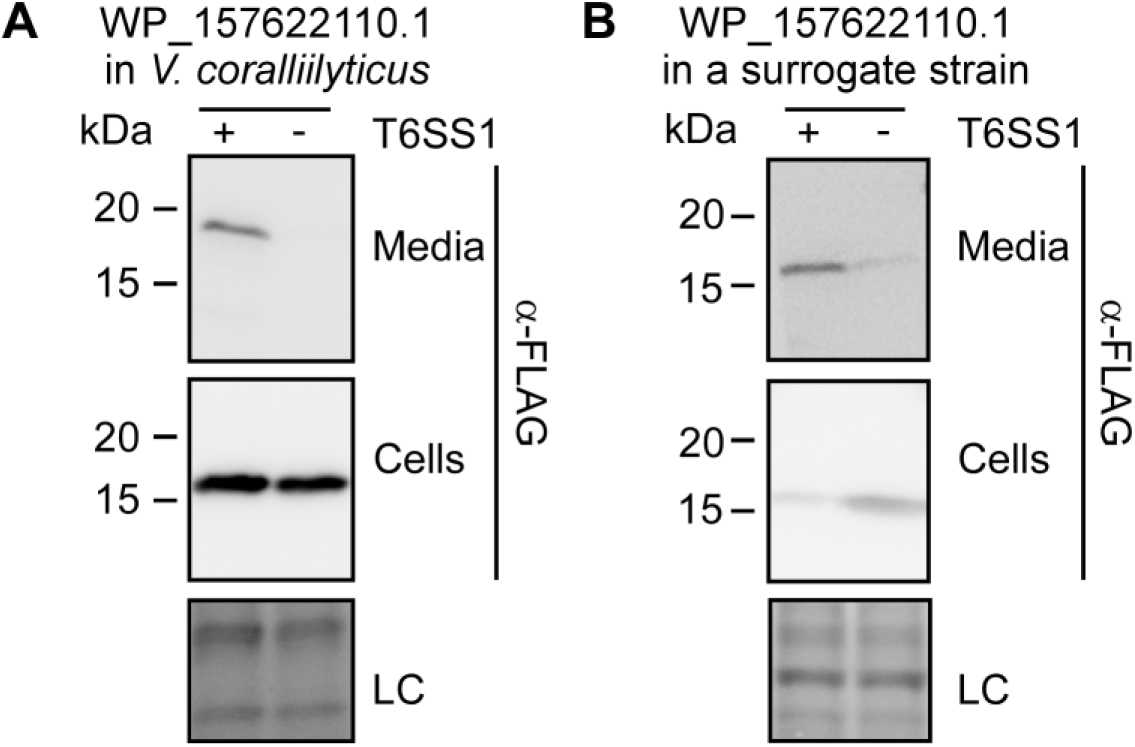
RIX-containing proteins are secreted by their endogenous T6SS. Expression (cells) and secretion (media) of a C-terminal FLAG-tagged WP_157622110.1 expressed from a pBAD33.1^F^-based plasmid in **(A)** a *V. coralliilyticus* BAA-450 wild-type strain (T6SS1^+^) or its ∆*hcp1* derivative (T6SS1^−^), or **(B)** a *V. parahaemolyticus* RIMD 2210633-derivative surrogate (T6SS1^+^) strain or its ∆*hcp1* derivative (T6SS1^−^). Samples were grown in MLB supplemented with chloramphenicol and 0.05% (wt/vol) L-arabinose (to induce expression from the plasmid) for 4 h at 30°C. Loading control (LC) is shown for the total protein lysates.

### A RIX tether-dependent mechanism for T6SS effector delivery

Unlike most RIX domain-containing proteins, which have a predicted C-terminal toxin domain, WP_157622110.1 has a predicted C-terminal TTR (Transthyretin-like) domain (TTR_1; see **Table 1**). This domain was previously identified at C-terminal extensions of secreted T6SS spike components, such as VgrG and PAAR, and it was suggested to function as an internal adaptor for cargo effectors ^12,39–41^. Therefore, we reasoned that WP_157622110.1 is not a toxic effector.

In *V. coralliillyticus* BAA-450, WP_157622110.1 is encoded by the first gene in a three-gene operon; notably, homologous modules comprising the TTR domain and the two downstream encoded proteins are also found fused to the T6SS-secreted components PAAR and VgrG in other bacteria (**Fig. 5A**). The genetic composition of the three-gene operon and the presence of the TTR domain in WP_157622110.1 led us to hypothesize that: (1) the second and third genes in the operon encode an antibacterial T6SS cargo effector and its cognate immunity protein, and (2) the RIX domain-containing protein acts as a tether that mediates the effector’s delivery via T6SS.

**Fig. 5.**
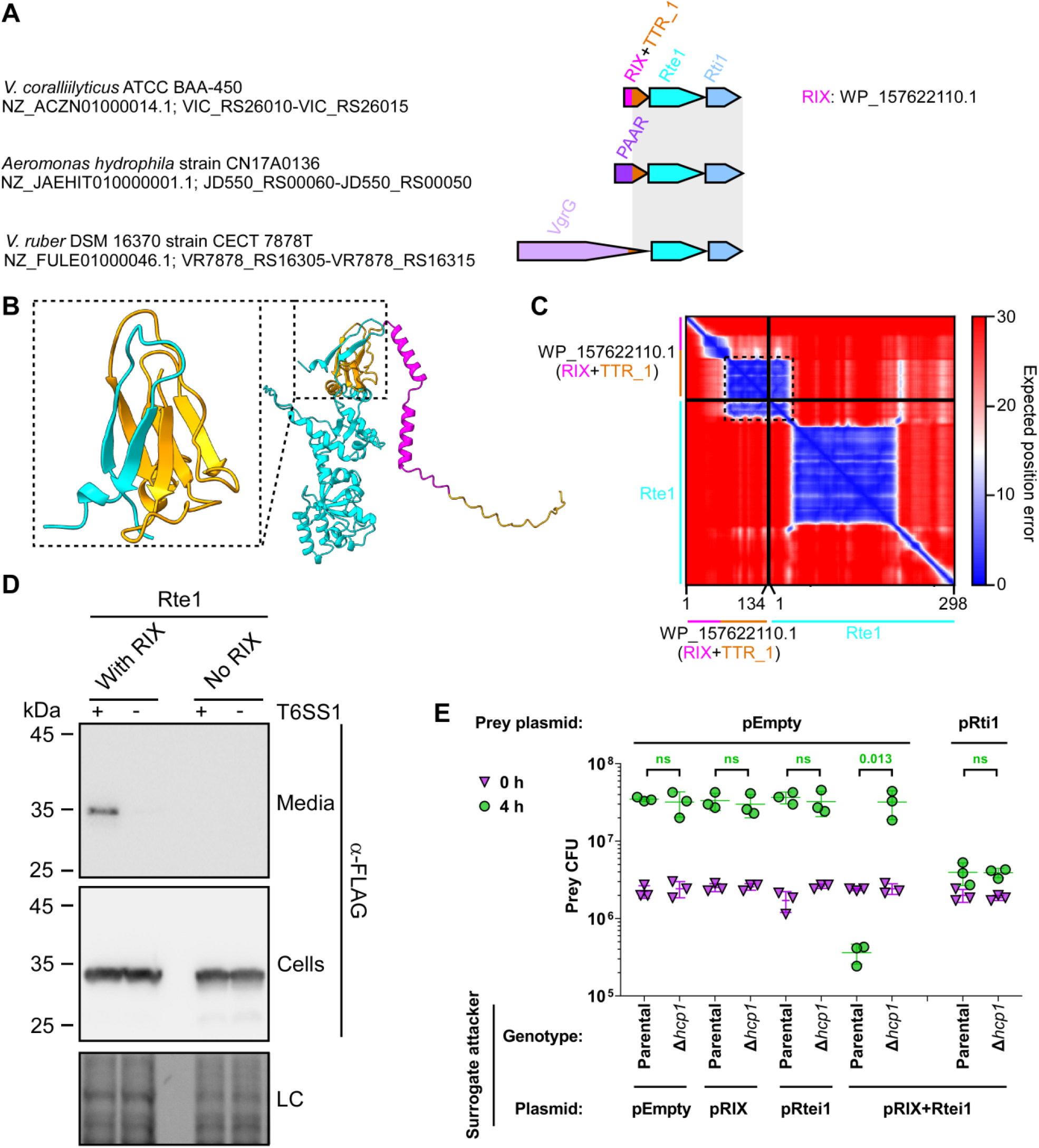
RIX-containing proteins can serve as tether for T6SS cargo effectors. **(A)** The gene structure of the operon encoding WP_157622110.1 in *V. coralliilyticus* BAA-450. Operons encoding a module homologous to the C-terminal extension of WP_157622110.1, Rte1, and Rti1 are shown below; a gray rectangle denotes the region of homology. The strain names, the GenBank accession numbers, and the locus tags are provided. Genes are denoted by arrows indicating the direction of transcription. The names of encoded proteins or domains are denoted above. **(B)** An AlphaFold2 structure prediction of the complex between WP_157622110.1 (orange, the region corresponding to RIX is shown in magenta) and Rte1 (cyan). The interaction interface, corresponding to amino acids 73-134 of WP_157622110.1 (orange) and 1-25 of Rte1 (cyan), is shown inside the dashed rectangle on the left. **(C)** The predicted aligned error of the complex shown in (B). A low predicted aligned error value indicates that the predicted relative position and orientation of two residues is well defined. **(D)** Expression (cells) and secretion (media) of the C-terminal FLAG-tagged Rte1^H94A^, either encoded alone (No RIX) or with the upstream-encoded WP_157622110.1 (With RIX), expressed from pBAD33.1^F^-based plasmids in a *V. parahaemolyticus* RIMD 2210633-derivative surrogate strain (T6SS1^+^) or its ∆*hcp1* derivative strain (T6SS1^−^). Samples were grown in MLB supplemented with chloramphenicol and 0.01% (wt/vol) L-arabinose (to induce expression from plasmids) for 4 h at 30°C. Loading control (LC) is shown for the total protein lysates. **(E)** Viability counts (CFU) of *V. parahaemolyticus* RIMD 2210633 ∆*hcp1* prey strains harboring either an empty plasmid (pEmpty) or a plasmid for the arabinose-inducible expression of Rti1 (pRti1) before (0 h) and after (4 h) co-incubation with a surrogate attacker strain (*V. parahaemolyticus* RIMD 2210633 derivative) or its T6SS1^−^ derivative (∆*hcp1*) carrying an empty plasmid or a plasmid for the arabinose-inducible expression of WP_157622110.1 (pRIX), Rte1 and Rti1 (pRtei1), or the three-gene operon including WP_157622110.1, Rte1 and Rti1 (pRIX-Rtei1). The statistical significance between samples at the 4 h timepoint was calculated using an unpaired, two-tailed Student’s *t*-test; ns, no significant difference (p>0.05). Data are shown as the mean ± SD; *n* = 3.

Before addressing these hypotheses experimentally, we used AlphaFold2 to predict whether the RIX-containing protein interacts with its downstream encoded protein. Indeed, AlphaFold2 predicted that the C-terminus of the RIX-containing WP_157622110.1, corresponding to the TTR domain, interacts with the N-terminus of the predicted effector (WP_006958655.1; hereafter named Rte1 for <u>R</u>IX-tethered <u>e</u>ffector <u>1</u>) to complete a barrel-like fold (**Fig. 5B**). This inter-domain prediction was made with a high degree of certainty, according to the AlphaFold2 predicted aligned error analysis (**Fig. 5C**). In addition, an AlphaFold2 structure prediction indicated that Rte1 and its predicted immunity protein, WP_0507786602.1 (hereafter named Rti1 for <u>R</u>IX-<u>t</u>ethered <u>i</u>mmunity <u>1</u>), interact with a high degree of certainty (**Supplementary Fig. S7A**,**B**). Furthermore, sequence (HHpred ^42^) and structure prediction (AlphaFold2 prediction followed by DALI server analysis ^43^) analyses suggest that Rte1 contains an ADPRT-like domain with a fold similar to that of the pertussis toxin (**Supplementary Fig. S7A**), to which Rti1 is expected to bind (**Supplementary Fig. S7A**,**B**), and a C-terminal glycine-zipper-like domain. A conservation logo assembled from a multiple sequence alignment of Rte1 homologs revealed conserved residues (H94, S106, and E174) that are similar to the catalytic residues of ADPRT toxins ^44^, which localize to a cleft within the predicted Rte1 structure (**Supplementary Fig. S7A**,**C**); this cleft corresponds to the ADPRT-like domain active site and is predicted to be occluded by Rti1 (**Supplementary Fig. S7A**).

Using these predictions, we set out to test our hypotheses. First, we determined whether Rte1 is toxic when expressed in bacteria. Indeed, its expression in *E. coli* was toxic; however, it was antagonized by co-expressing Rti1 (**Supplementary Fig. S7D**). Substitution of either H94 or S106, corresponding to conserved residues that were identified as the possible active site in the sequence and the structural analyses of Rte1, for alanine abolished the toxic activity of Rte1 in *E. coli* (**Supplementary Fig. S7E**). The expression of all Rte1 forms in *E. coli* was confirmed in immunoblots (**Supplementary Fig. S7F**).

Next, we tested whether Rte1 is secreted in a T6SS-dependent manner from the T6SS surrogate system, in the presence and absence of its cognate RIX domain-containing protein. To avoid self-intoxication of the surrogate strain by the effector, we used an inactive mutant of Rte1 (Rte1^H94A^; see **Supplementary Fig. S7E**). As shown in **Fig. 5D**, a plasmid-encoded Rte1^H94A^ was secreted in a T6SS-dependent manner when the upstream-encoded RIX domain-containing protein, WP_157622110.1, was co-expressed; however, in the absence of the RIX-domain-containing protein, Rte1^H94A^ was not detected in the medium. These results indicate that the RIX and TTR domains-containing protein is required for T6SS-mediated secretion of Rte1.

In addition, we investigated whether Rte1 and Rti1 function as an antibacterial T6SS effector and immunity pair in which the effector is dependent on a tether protein for delivery. To this end, we performed self-competition assays using the surrogate T6SS system. As shown in **Fig. 5E**, T6SS-dependent toxicity against a sensitive prey strain was only observed when Rte1 and Rti1 were expressed together with the upstream-encoded RIX domain-containing protein (pRIX+Rtei1), but not when they were expressed alone (pRtei1). Notably, expression of the RIX domain-containing protein alone (pRIX) did not result in prey intoxication. Expression of Rti1 from a plasmid in the prey strain protected it from the T6SS-mediated toxicity. Taken together, our results support a role for a RIX domain-containing protein as a tether that mediates the delivery of a cargo effector via T6SS.

When we examined other RIX domain-containing proteins that had no predicted toxin domain at their C-terminus, we identified two additional types of predicted RIX-associated T6SS tethers. As exemplified by WP_095560627.1 and WP_152436132.1, these RIX domain-containing proteins are encoded by the first gene in a three-gene operon (**Supplementary Fig. S8A**). Notably, the effectors encoded by the second gene within these two operons are known: a Tae4 peptidoglycan amidase effector ^17^and a homolog of the TseH cysteine hydrolase effector ^41,45^, respectively. The third gene in these operons encodes the predicted cognate immunity protein. Similar to the Rte1 tether example described above, a module encompassing the C-terminal extension of the RIX domain (i.e., the predicted loading platform), the effector, and the immunity protein can be found fused to a PAAR domain in other vibrios. Structure predictions using AlphaFold2 suggest that the RIX-fused C-terminal domain and the N-terminus of the downstream encoded effector interact (**Supplementary Fig. S8B**), thus further supporting the notion that RIX domain-containing proteins can serve as T6SS tethers for cargo effectors.

## Discussion

In this work, we identified a new class of T6SS-secreted proteins that are widespread in *Vibrionaceae*. These proteins share an N-terminal domain that we named RIX; however, they have polymorphic C-terminal extensions with predicted antibacterial, anti-eukaryotic, or cargo binding activities (i.e., tethers). We showed that RIX is necessary for T6SS-mediated secretion, and we experimentally confirmed the T6SS-mediated secretion of six representative RIX-containing proteins: two antibacterial effectors (Tme1 and WP_005534959.1), three anti-eukaryotic effectors (WP_005536620.1, WP_005530005.1, and WP_038863399.1), and one tether (WP_157622110.1). In addition, we identified a novel, non-RIX-containing T6SS cargo effector, Rte1. Notably, without RIX, the identification of these polymorphic T6SS effectors would not have been trivial, since all RIX-encoding genes are orphan (i.e., they are not located near a T6SS-associated gene), and because many of the predicted toxin domains found in RIX-containing proteins have not been previously associated with T6SS effectors.

Revealing many RIX-containing proteins that have predicted anti-eukaryotic toxin domains suggests that we have underappreciated the potential role played by T6SS in the interactions between vibrios and eukaryotic organisms, whether they are hosts or predators. Although T6SS was originally described as an anti-eukaryotic determinant, allowing *V. cholerae* to escape predation by grazing amoeba ^1^, most *Vibrio* T6SSs investigated to date were shown to play a role in interbacterial competitions ^46–52^, and only a handful of *Vibrio* T6SSs have been implicated in anti-eukaryotic activities. These *Vibrio* T6SSs include T6SS1 and T6SS3 in *V. proteolyticus* ^48,53^, T6SS1 in *V. tasmanienses* ^54^, and T6SS in *V. crassostreae* ^55^. Several additional *Vibrio* T6SSs were also suggested to play a role in interactions with eukaryotes based on the presence of a predicted anti-eukaryotic MIX-effector in their genome ^23^. Furthermore, only a few T6SS effectors with anti-eukaryotic activities have been described to date, in vibrios and in other bacteria ^5^. Therefore, our identification of diverse families of RIX-containing effectors with known or predicted anti-eukaryotic activities substantially enlarges this list, and it may instigate future studies to identify the eukaryotic targets and the mechanisms of action of these novel effectors. Future work is also required to identify the potential eukaryotic organisms that are targeted by these potentially anti-eukaryotic T6SSs found in vibrios, and to determine whether their role is offensive or defensive.

In addition to effectors, RIX domains are also found in T6SS tethers, revealing a new mechanism of tether-mediated secretion for T6SS cargo effectors. Our results indicated that RIX domains replace PAAR and VgrG proteins, which are secreted structural components of the T6SS, in modules comprising a C-terminal loading platform extension and a cargo effector and immunity pair. We experimentally demonstrated that a new cargo effector, Rte1, required an upstream-encoded RIX-containing protein in order to be secreted by the T6SS and delivered into prey cells during bacterial competition. This tether-mediated secretion mechanism is similar to the recently described mechanism involving a MIX domain-containing co-effector ^22^, yet it is distinct since the RIX-containing tether is secreted via T6SS on its own, whereas the MIX-containing co-effector requires its effector partner for secretion. Notably, our attempts to determine the T6SS tube-spike component with which RIX domain-containing proteins interact have been inconclusive; therefore, future work is also required to determine the mechanism governing the secretion of RIX domain-containing proteins.

In conclusion, we identified a new class of polymorphic T6SS cargo effectors widespread in vibrios, and we revealed a new, tether-mediated secretion mechanism for T6SS cargo effectors. In future work, we will determine how RIX-containing proteins are loaded onto the secreted T6SS, and we will explore the potential use of RIX domains and RIX-containing tethers to deliver engineered, non-canonical cargo effectors via T6SS to be used as bio-treatments ^56^ or molecular biology tools ^57^.

## Materials and Methods

### Strains and Media

For a complete list of strains used in this study, see **Supplementary Table S1**. *Escherichia coli* strains BL21 (DE3), DH5α (λ-pir), and *Pseudomonas aeruginosa* PAO1 were grown in 2xYT broth (1.6% wt/vol tryptone, 1% wt/vol yeast extract, and 0.5% wt/vol NaCl) or on lysogeny broth agar (LB; 1.5% wt/vol) plates at 37°C, or at 30°C when harboring effector expression plasmids. The media were supplemented with chloramphenicol (10 μg/ml) and/or kanamycin (30 μg/ml) to maintain plasmids and with 0.4% (wt/vol) glucose to repress protein expression from the arabinose-inducible promoter, P*bad*. To induce expression from P*bad*, L-arabinose was added to the media at 0.05-0.1% (wt/vol), as indicated.

*Vibrio parahaemolyticus* strains BB22OP, RIMD 2210633, and their derivatives, as well as *Vibrio natriegens* ATCC 14048 and *Vibrio coralliilyticus* ATCC BAA-450, were grown in Marine Lysogeny Broth (MLB; LB containing 3% w/v NaCl) and on Marine Minimal Media (MMM) agar plates (1.5% wt/vol agar, 2% wt/vol NaCl, 0.4% wt/vol galactose, 5 mM MgSO_4_, 7 mM K_2_SO_4_, 77 mM K_2_HPO_4_, 35 mM KH_2_PO_4_, and 2 mM NH_4_Cl) at 30°C. The media were supplemented with chloramphenicol (10 μg/ml) or kanamycin (250 μg/ml) to maintain plasmids. To induce expression from P*bad*, L-arabinose was added to media at 0.01-0.1% (wt/vol), as indicated.

*Saccharomyces cerevisiae* BY4741 (MATa, his3Δ0, leu2Δ0, met15Δ0, and ura3Δ0) yeast were grown in Yeast Extract–Peptone–Dextrose broth (YPD; 1% wt/vol yeast extract, 2% wt/vol peptone, and 2% wt/vol glucose) or on YPD agar (2% wt/vol) plates at 30°C. Yeast containing plasmids that provide prototrophy to leucine were grown in Synthetic Drop-out media (SD; 6.7 g/l yeast nitrogen base without amino acids, 1.4 g/l yeast synthetic drop-out medium supplement) supplemented with histidine (2 ml/l from a 1% wt/vol stock solution), tryptophan (2 ml/l from 1% wt/vol stock solution), uracil (10 ml/l from a 0.2% wt/vol stock solution), and glucose (4% wt/vol). For galactose-inducible expression from a plasmid, cells were grown in SD media or on SD agar plates supplemented with galactose (2% wt/vol) and raffinose (1% wt/vol).

### Plasmid construction

For a complete list of plasmids used in this study, see **Supplementary Table S2**. For expression in bacteria or yeast, the coding sequences (CDS) of the following protein accession numbers: WP_015297525.1 (Tme1), WP_003088027.1 (Tse1), WP_005534959.1, WP_005534960.1, WP_005536620.1, WP_005530005.1, WP_038863399.1, WP_157622110.1, WP_006958655.1, and WP_050778602.1 were PCR amplified from the respective genomic DNA of the encoding bacterium. Next, amplicons were inserted into the multiple cloning site (MCS) of pBAD^K^/Myc-His, pBAD33.1^F^, or their derivatives using the Gibson assembly method ^58^. Plasmids were introduced into *E. coli* BL21 (DE3) or DH5α (λ-pir) by electroporation, and into vibrios via conjugation. Transconjugants were selected on MMM agar plates supplemented with the appropriate antibiotics to select clones containing the desired plasmids. For galactose-inducible expression in yeast, genes were inserted into the MCS of the shuttle vector pGML10 (Riken) using the Gibson assembly method, in-frame with a C-terminal Myc-tag. Yeast transformations were performed using the lithium acetate method, as previously described ^59^.

### Construction of deletion strains

The construction of the *V. parahaemolyticus* RIMD 2210633 surrogate strain and of the *V. parahaemolyticus* BB22OP and *V. coralliilyticus* BAA-450 derivatives was reported previously ^29,37^. Briefly, 1 kb sequences upstream and downstream of each gene to be deleted were cloned into pDM4, a Cm^R^OriR6K suicide plasmid. The pDM4 constructs were transformed into *E. coli* DH5α (λ pir) by electroporation, and then transferred into *Vibrio* isolates via conjugation. Transconjugants were selected on agar plates supplemented with chloramphenicol, and then counter-selected on agar plates containing 15% (wt/vol) sucrose for loss of the sacB-containing plasmid. Deletions were confirmed by PCR.

### Toxicity assays in *E. coli*

For testing the toxicity of proteins during the growth of bacteria in suspension, *E. coli* strains carrying arabinose-inducible expression plasmids were grown in 2xYT broth supplemented with the appropriate antibiotics and 0.4% (wt/vol) glucose (to repress expression from the P*bad* promotor) at 30°C. Overnight cultures were washed twice with fresh 2xYT broth, normalized to an OD_600_ of 0.01 in 2xYT broth and then transferred to 96-well plates (200 µl per well) in quadruplicate. Cultures were grown at 37°C in a BioTek SYNERGY H1 microplate reader with constant shaking (205 cpm). After 2 h, L-arabinose was added to a final concentration of 0.05% (wt/vol) to induce protein expression. OD_600_ readings were acquired every 10 min.

For testing the toxicity of proteins during the growth of bacteria on solid media, *E. coli* strains carrying arabinose-inducible expression plasmids were streaked onto LB agar plates supplemented with the appropriate antibiotics and 0.4% (wt/vol) glucose (repressing plates) or 0.1% (wt/vol) L-arabinose (inducing plates). Plates were incubated for 16 h at 37°C. The experiments were performed at least three times with similar results. Results from a representative experiment are shown.

### Protein expression in *E. coli*

*E. coli* cultures harboring arabinose-inducible expression plasmids were grown overnight in 2xYT broth supplemented with appropriate antibiotics to maintain the expression plasmids. Bacterial cultures were then washed with fresh 2xYT and resuspended in 3 ml of 2xYT supplemented with appropriate antibiotics. Next, bacterial cultures were incubated with constant shaking (220 rpm) at 37°C for 2 hours. After 2 h, L-arabinose was added to a final concentration of 0.05% (wt/vol) to induce protein expression, and cultures were grown for 2 additional hours. Cells equivalent to 0.5 OD_600_ units were collected, and cell pellets were resuspended in 50 µl of 2x Tris-glycine SDS sample buffer (Novex, Life Sciences). Next, samples were boiled at 95°C for 10 minutes, and then loaded onto TGX stain-free gels (Bio-Rad) for SDS-PAGE. Proteins were transferred onto nitrocellulose membranes, which were immunoblotted with α-FLAG (Sigma-Aldrich, F1804), α-Myc (Santa Cruz, 9E10, sc-40), or custom-made α-VgrG1 ^60^ antibodies at 1:1000 dilution, as indicated. Finally, protein signals were visualized in a Fusion FX6 imaging system (Vilber Lourmat) using enhanced chemiluminescence (ECL). The experiments were performed at least three times with similar results. Results from a representative experiment are shown.

### Toxicity assays in yeast

The experiments were performed as previously described ^59^. Briefly, yeast cells were grown overnight in appropriate media supplemented with 4% (wt/vol) glucose, washed twice with ultrapure milli-Q water, and then normalized to an OD_600_ of 1.0 in milli-Q water. Next, 10-fold serial dilutions were spotted onto synthetic dropout agar plates containing either 4% (wt/vol) glucose (repressing plates) or 2% (wt/vol) galactose and 1% (wt/vol) raffinose (inducing plates). The plates were incubated at 30°C for two days. The experiments were performed at least three times with similar results. Results from a representative experiment are shown.

### Protein expression in yeast

Yeast cells harboring galactose-inducible expression plasmids were grown overnight in selective media supplemented with glucose, washed twice with ultrapure milli-Q water, and then normalized to OD_600_ = 1.0 in selective media supplemented with galactose and raffinose to induce protein expression. Next, the cells were grown at 30°C for 16 h, and 1.0 OD_600_ units of cells were pelleted and lysed, as previously described ^59^. Proteins were detected in immunoblots using α-Myc antibodies at a 1:1000 dilution. The experiments were performed at least three times with similar results. Results from a representative experiment are shown.

### Bacterial competition assays

Competition assays were performed as previously described ^46^, with minor modifications. Briefly, attacker and prey *Vibrio* strains were grown overnight, normalized to an OD_600_ of 0.5, and then mixed at a 4:1 (attacker:prey) ratio in triplicate. Then, 25 μl of the mixtures were spotted onto MLB agar competition plates supplemented with 0.05% (wt/vol) L-arabinose when induction of protein expression from a plasmid was required. Competition plates were incubated at the indicated temperatures for 4 h. The colony-forming units (CFU) of the prey strains at t = 0 h were determined by plating 10-fold serial dilutions on selective media plates. After 4 h of co-incubation of the attacker and prey mixtures on the competition plates, the bacteria were harvested and the CFUs of the surviving prey strains were determined by plating 10-fold serial dilutions on selective media plates. The experiments were performed at least three times with similar results. Results from a representative experiment are shown.

### Protein secretion assays

Secretion assays were performed as previously described ^22^, with minor modifications. Briefly, *Vibrio* strains were grown overnight and then normalized to an OD_600_ of 0.18 in 3 ml MLB broth supplemented with appropriate antibiotics and 0.01-0.05% (wt/vol) L-arabinose, when expression from an arabinose-inducible plasmid was required. Bacterial cultures were incubated with constant shaking (220 rpm) at the indicated temperatures for the specified durations. For expression fractions (cells), cells equivalent to 0.5 OD_600_ units were collected, and cell pellets were resuspended in 50 µl of 2x Tris-glycine SDS sample buffer (Novex, Life Sciences). For secretion fractions (media), supernatant volumes equivalent to 10 OD_600_ units were filtered (0.22 µm), and proteins were precipitated using the deoxycholate and trichloroacetic acid method ^61^. The precipitated proteins were washed twice with cold acetone, and then air-dried before resuspension in 20 µl of 100 mM Tris-Cl (pH = 8.0) and 20 µl of 2X protein sample buffer with 5% β-mercaptoethanol. Next, samples were incubated at 95°C for 5 or 10 min and then resolved on TGX Stain-free gel (Bio-Rad). The proteins were transferred onto 0.2 µm nitrocellulose membranes using Trans-Blot Turbo Transfer (Bio-Rad) according to the manufacturer’s protocol. Membranes were then immunoblotted with α-FLAG (Sigma-Aldrich, F1804) or custom-made α-VgrG1 ^60^ antibodies at 1:1000 dilution. Protein signals were visualized in a Fusion FX6 imaging system (Vilber Lourmat) using enhanced chemiluminescence (ECL) reagents. The experiments were performed at least three times with similar results. Results from a representative experiment are shown.

### Identifying RIX-containing proteins

The position-specific scoring matrix (PSSM) of RIX was constructed using the N-terminal 55 residues of Tme1 (WP_015297525.1) from *V. parahaemolyticus* BB22OP. Five iterations of PSI-BLAST were performed against the RefSeq protein database. In each iteration, a maximum of 500 hits with an expect value threshold of 10^−6^ were used. Compositional adjustments and filters were turned off. The genomic neighborhoods of RIX-containing proteins (**Supplementary Dataset S1**) were analyzed as described previously ^23,29^. Duplicated protein accessions appearing in the same genome in more than one genomic accession were removed if the same downstream protein existed at the same distance. The T6SS core components in the RIX-containing bacterial genomes (**Supplementary Dataset S2**) were identified as previously described ^16^).

### Illustration of the conserved residues of the RIX domain

RIX domain sequences were aligned using Clustal Omega ^62^. Aligned columns not found in the RIX domain of Tme1 were discarded. The RIX domain-conserved residues were illustrated using the WebLogo server ^63^ (https://weblogo.threeplusone.com).

### Constructing a phylogenetic tree of RIX-encoding bacterial strains

DNA sequences of *rpoB* were aligned using MAFFT v7.505 FFT-NS-2 (https://mafft.cbrc.jp/alignment/server) ^64^. Partial and pseudogene sequences were discarded. The evolutionary history was inferred using the neighbor-joining method ^65^ with the Jukes-Cantor substitution model (JC69). The analysis included 686 nucleotide sequences and 4,021 conserved sites.

### Protein structure predictions

Predicted protein structures were downloaded from the AlphaFold Protein Structure Database ^36,66^ in August 2022 (https://alphafold.ebi.ac.uk/). The structures of proteins that were not available on the database and of protein complexes were predicted in ColabFold: AlphaFold2 using MMseqs2 ^35^. All PDB files used in this work are available as **Supplementary File S1**. Protein structures were visualized using ChimeraX 1.4 ^67^.

### Analyses of RIX C-terminal sequences

Amino acid sequences C-terminal to RIX were clustered in two dimensions using CLANS ^34^. To predict the activities or domains in each cluster, at least two representative sequences (when more than one was available) were analyzed using the NCBI Conserved Domain Database ^68^ and HHpred ^42^. If no activity or domain could be predicted, the protein sequences were further used for AlphaFold2 structure prediction ^35^ followed by a 3D protein structure comparison in the Dali server ^43^.

### Illustration of the conserved residues of Rte1

Homologs of Rte1 (WP_006958655.1) from *Vibrio coralliilyticus* were identified using PSI-BLAST (4 iterations; a maximum of 500 hits with an expect value threshold of 10^−6^ and a query coverage of 70% were used). Rte1 homologs were aligned using Clustal Omega and conserved residues were illustrated using WebLogo 3.

## Acknowledgments

This project received funding from the European Research Council under the European Union’s Horizon 2020 research and innovation program (grant agreement no. 714224), and from the Israel Science Foundation (grant no. 920/17 to D Salomon, and grant no. 1362/21 to D Salomon and E Bosis). CM Fridman was supported by a scholarship from the Clore Israel Foundation and by a scholarship for outstanding doctoral students from the Orthodox community from the Council for Higher Education. We thank Kinga Keppel and Biswanath Jana for technical assistance, and the rest of the members of the Salomon and Bosis labs for helpful discussions and suggestions. This work was performed in partial fulfillment of the requirements for a PhD degree for K Kanarek at the Sackler Faculty of Medicine, Tel Aviv University.

## Author Contributions

K Kanarek: conceptualization, investigation, methodology, and writing—original draft.

CM Fridman: investigation, methodology, and writing—review and editing.

E Bosis: conceptualization, investigation, methodology, funding acquisition, and writing—original draft.

D Salomon: conceptualization, supervision, funding acquisition, investigation, methodology, and writing—original draft.

## Conflict of Interest

The authors declare that they have no conflict of interest.

## Notes

### Competing Interest Statement

The authors have declared no competing interest.

