## Supplementary Information - Figures and Tables for "A new class of polymorphic T6SS effectors and tethers"


- **Supplementary Figures S1-S8**
- **Supplementary Datasets S1-S2**
- **Supplementary File S1**
- **Supplementary Tables S1-S2**
- **Supplementary References**

### Supplementary Figures

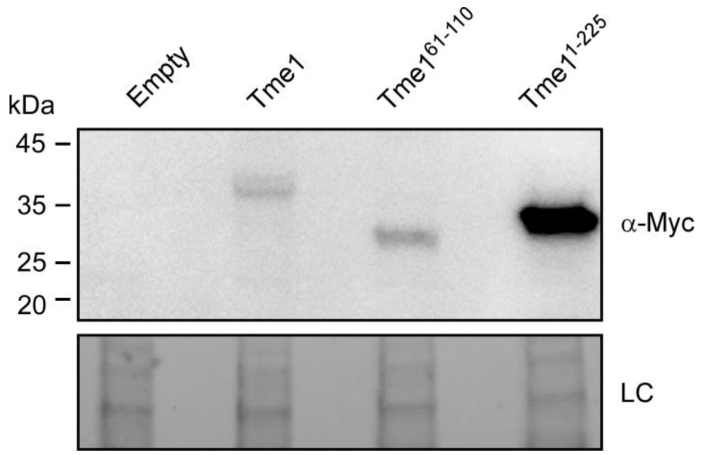


**Supplementary Fig. S1. Tme1 truncations are expressed in *E. coli*.** Expression of the indicated C-terminal Myc-His tagged Tme1 forms from arabinose-inducible pPER5-based plasmids in *E. coli* BL21 (DE3). Proteins were detected by immunoblotting using specific α-Myc antibodies. Loading control (LC) is shown for total protein lysates.

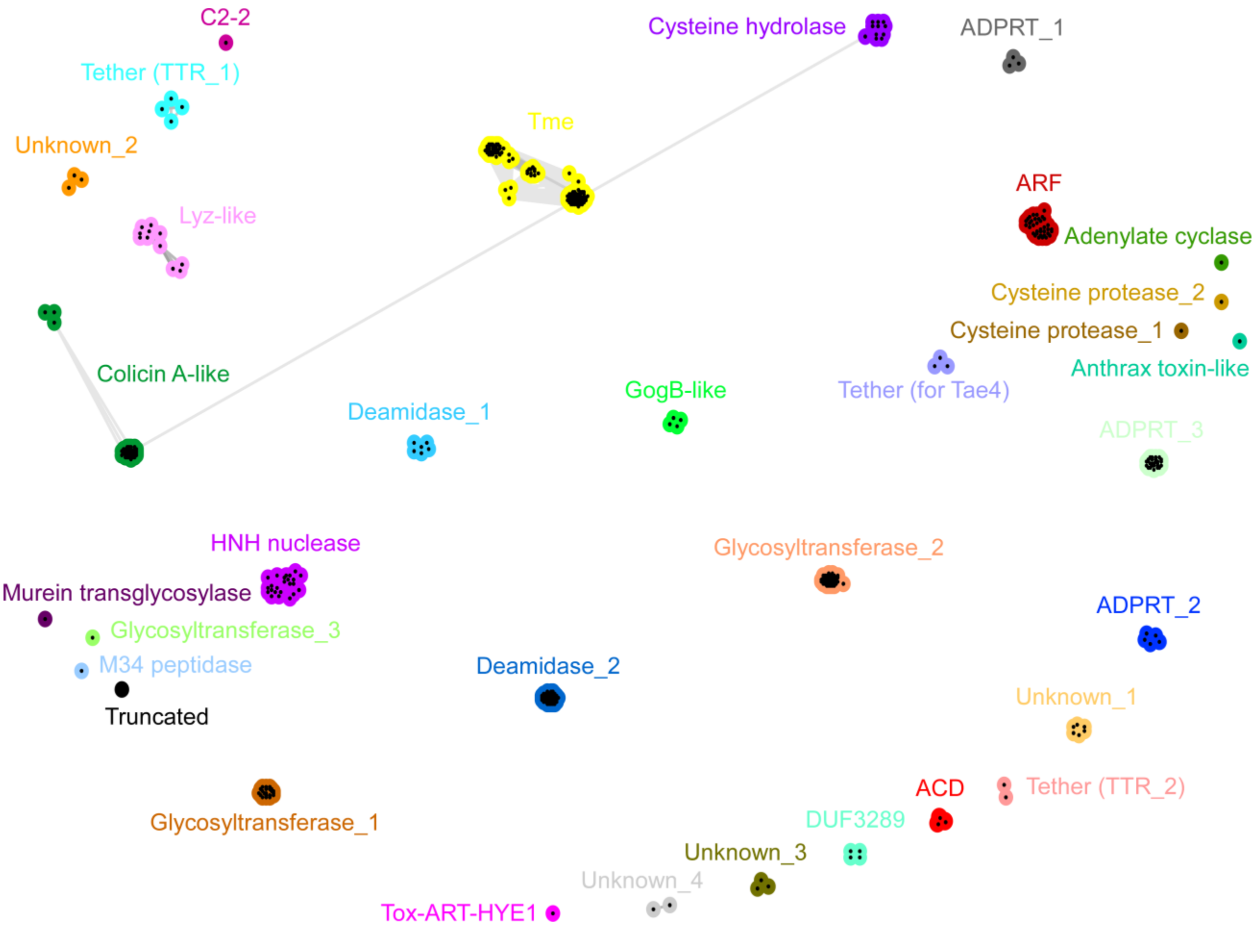


**Supplementary Fig. S2. RIX domain C-terminal extensions cluster into 33 distinct families.** C-termini of RIX-containing proteins clustered in two dimensions. Clustering was performed based on all-against-all sequence similarity, with nodes representing unique sequences and connecting lines representing the distances between sequences. The predicted activities or domains identified in each cluster are denoted.

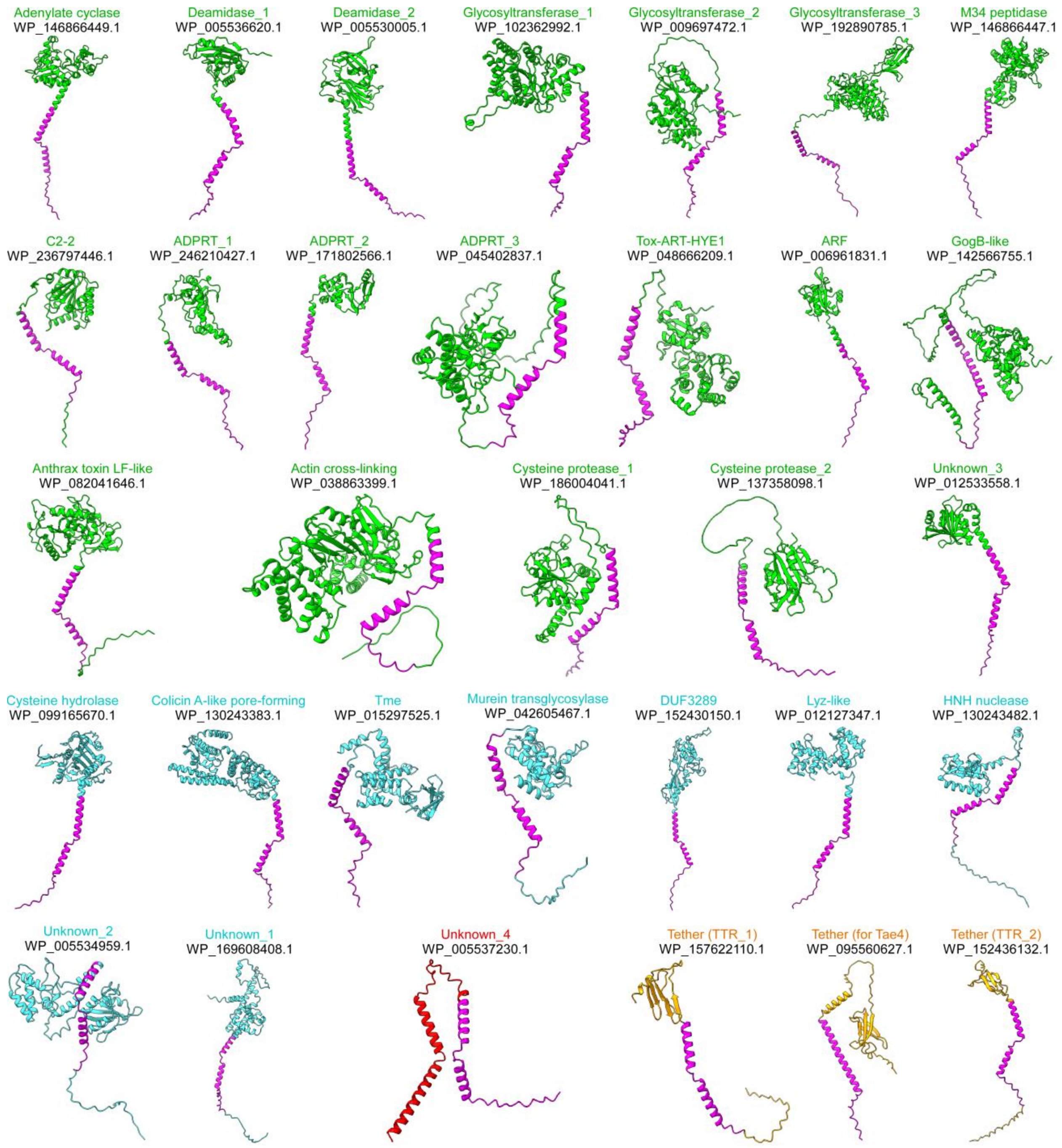


**Supplementary Fig. S3. AlphaFold2 structure predictions of RIX cluster representatives.** The structures of a representative protein from 32 RIX C-termini clusters (excluding the "truncated" cluster) were predicted using AlphaFold2. The protein accession number and the predicted activity or domain are denoted above. Regions corresponding to RIX are denoted in magenta; proteins with predicted anti-eukaryotic activities are denoted in green; proteins with predicted antibacterial activities are denoted in cyan; proteins with predicted tether activities are denoted in orange; C-terminal extensions with an unknown function are denoted in red.

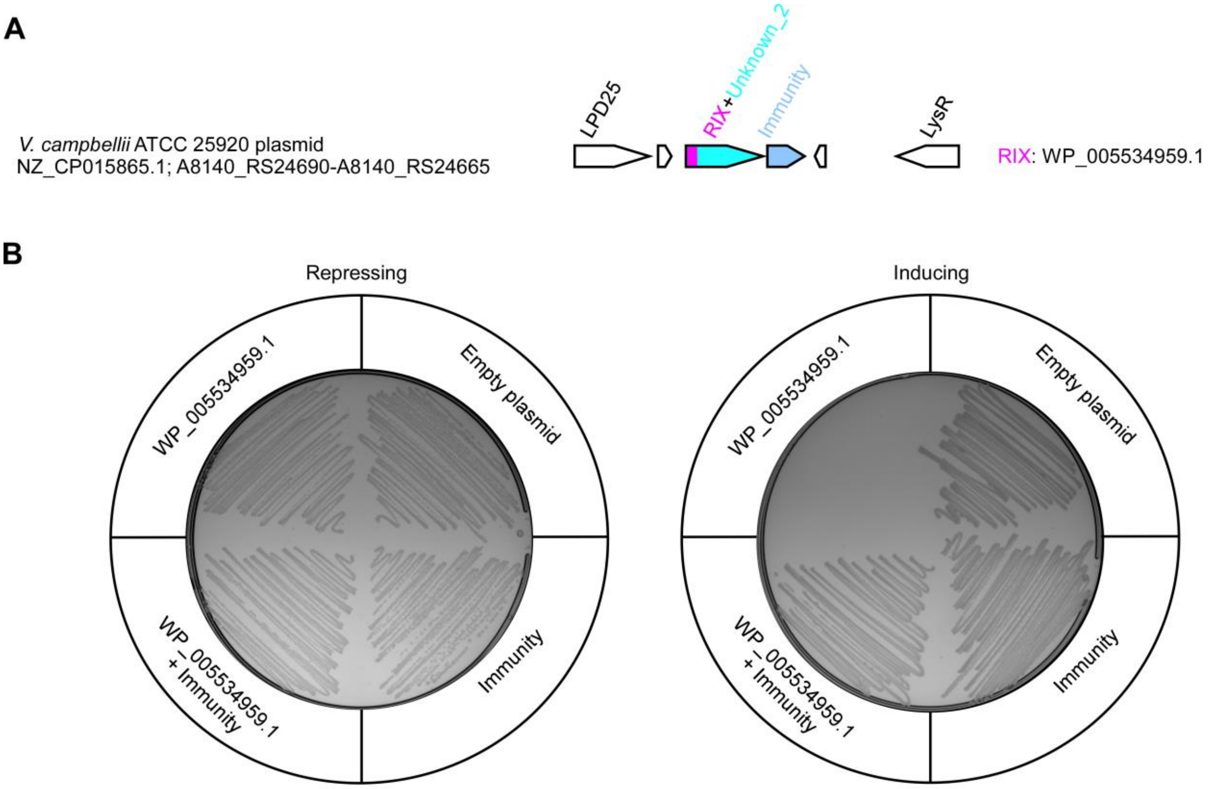


**Supplementary Fig. S4. The RIX-containing WP_005534959.1 is an antibacterial toxin. (A)** The genomic neighborhood of the gene encoding WP_005534959.1 in *V. campbellii* ATCC 25920. The strain name, the GenBank accession number, and the locus tags are provided. Genes are denoted by arrows indicating the direction of transcription. The names of encoded proteins or domains are denoted above. **(B)** Toxicity of WP_005534959.1 expressed from pBAD33.1$^{F}$-based arabinose-inducible plasmids in *E. coli* BL21 (DE3), with or without its downstream-encoded predicted immunity protein. Bacteria were streaked onto repressing (4% wt/vol glucose) or inducing (0.05% wt/vol arabinose) agar plates.

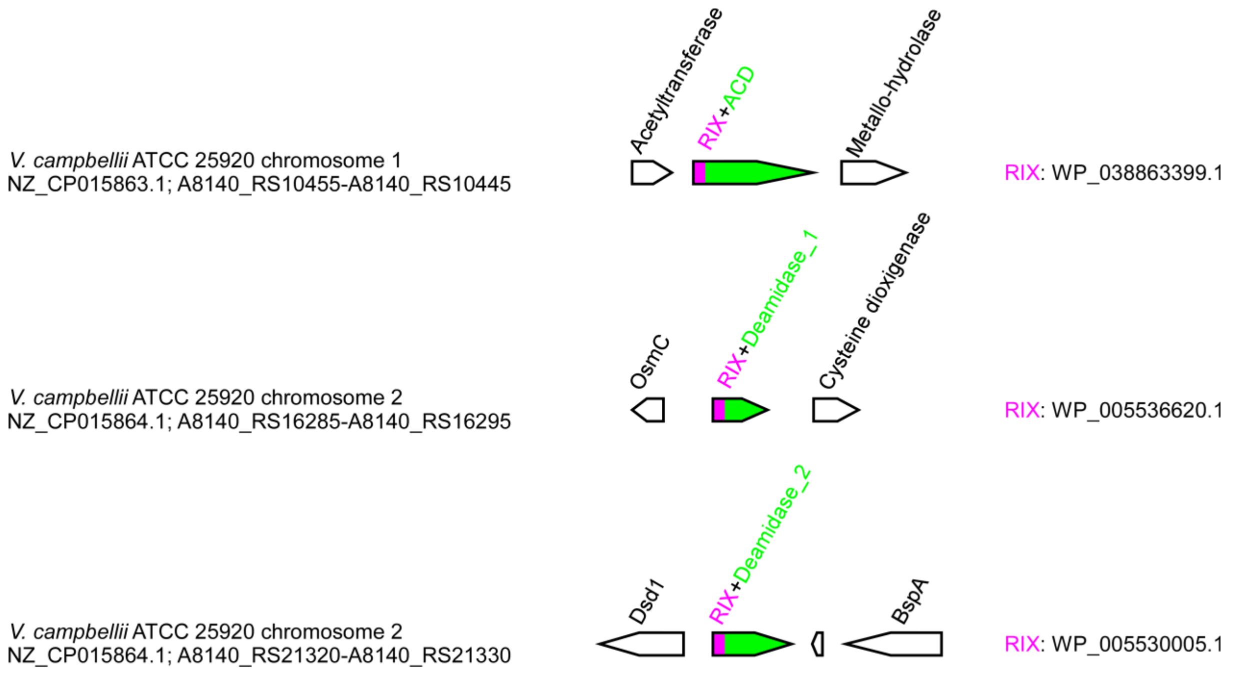


**Supplementary Fig. S5. Three genes encoding RIX-containing proteins with predicted anti-eukaryotic activities in *V. campbellii*.** The genomic neighborhood of the genes encoding the indicated RIX-containing proteins in *V. campbellii* ATCC 25920. The strain name, the GenBank accession number, and the locus tags are provided. Genes are denoted by arrows indicating the direction of transcription. The names of encoded proteins or domains are denoted above.

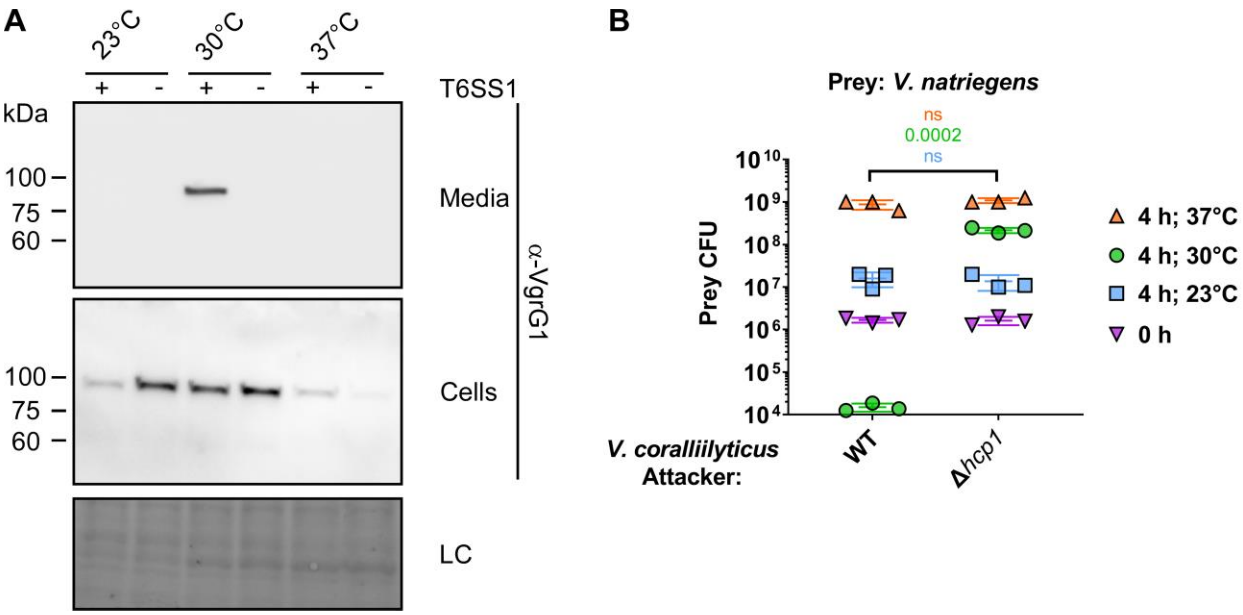


**Supplementary Fig. S6.** ***V. coralliilyticus*** **T6SS1 mediates antibacterial toxicity under warm, marine-like conditions. (A)** Expression (cells) and secretion (media) of VgrG1 from *V. coralliilyticus* BAA-450 wild type (T6SS1$^+$) or Δ*hcp1* (T6SS1$^-$) strains. Samples were grown in MLB media for 4 h at the indicated temperature. Loading control (LC) is shown for total protein lysates. **(B)** Viability counts of *V. natriegens* prey before (0 h) and after (4 h) co-incubation with the indicated *V. coralliilyticus* BAA-450 attacker at the indicated temperature. The statistical significance between samples at the 4 h timepoint (color coded to match the relevant samples) was calculated using an unpaired, two-tailed Student's *t*-test; ns, no significant difference (p>0.05). Data are shown as the mean ± SD; *n* = 3. WT, wild type.

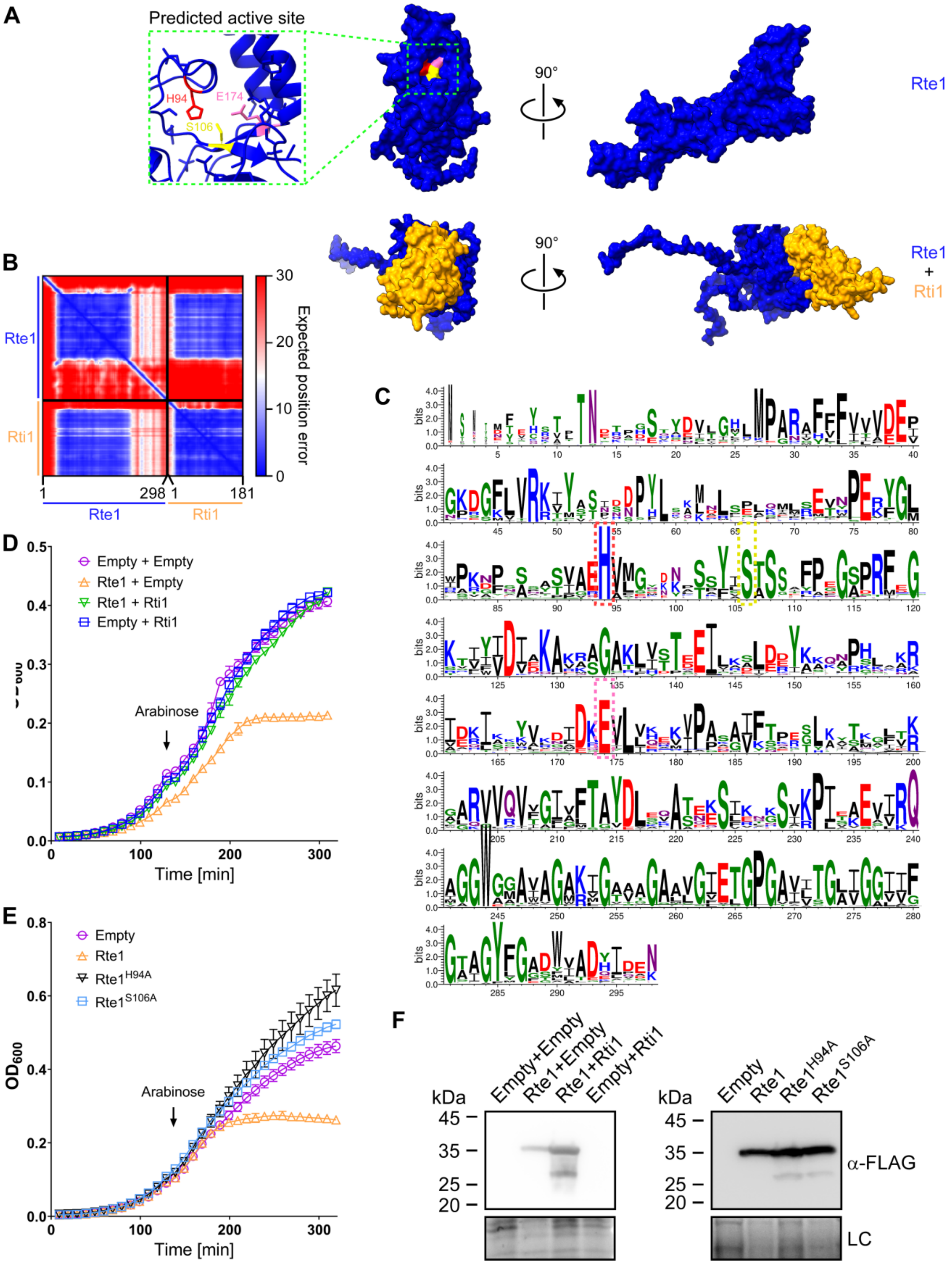

A
Predicted active site
H94
E174
S106
90°
Rte1
Rte1
+
Rti1
B
Rte1
Rti1
Expected position error
30
20
10
0
1
298
1
181
C
bits
D
Empty + Empty
Rte1 + Empty
Rte1 + Rti1
Empty + Rti1
Arabinose
Time [min]
E
Empty
Rte1
Rte1H94A
Rte1S106A
OD600
Arabinose
Time [min]
F
Empty+Empty
Rte1+Empty
Rte1+Rti1
Empty+Rti1
kDa
45
35
25
20
Empty
Rte1
Rte1H94A
Rte1S106A
α-FLAG
LC

**Supplementary Fig. S7. Rte1 and Rti1 are an antibacterial effector and immunity pair.** **(A)** The structure of Rte1, either alone (top) or in complex with Rti1 (bottom) was predicted using AlphaFold2. Predicted active site residues are shown inside the green dashed rectangle. **(B)** The predicted aligned error of the complex shown in (A). A low predicted aligned error value indicates that the predicted relative position and orientation of two residues is well defined. **(C)** A conservation logo of Rte1 is illustrated using WebLogo3, based on multiple sequence alignment of Rte1 homologs. The position numbers correspond to the amino acids in Rte1. Predicted active site residues are denoted by a dashed rectangle, and color coded to match the residues shown in (A). **(D-E)** Growth of *E. coli* BL21 (DE3) containing plasmids for the arabinose-inducible expression of the indicated proteins. An arrow denotes the timepoint at which arabinose (0.05% wt/vol) was added to the media. In (D), Rte1 and Rti1 were expressed from separate plasmids (pBAD33.1$^{F}$ and pBAD$^{K}$/Myc-His-based, respectively. **(F)** Expression of the indicated C-terminal FLAG tagged Rte1 forms from arabinose-inducible plasmids in *E. coli* BL21 (DE3) strains used in (D) and (E). Proteins were detected by immunoblotting using specific α-FLAG antibodies. Loading control (LC) is shown for total protein lysates.

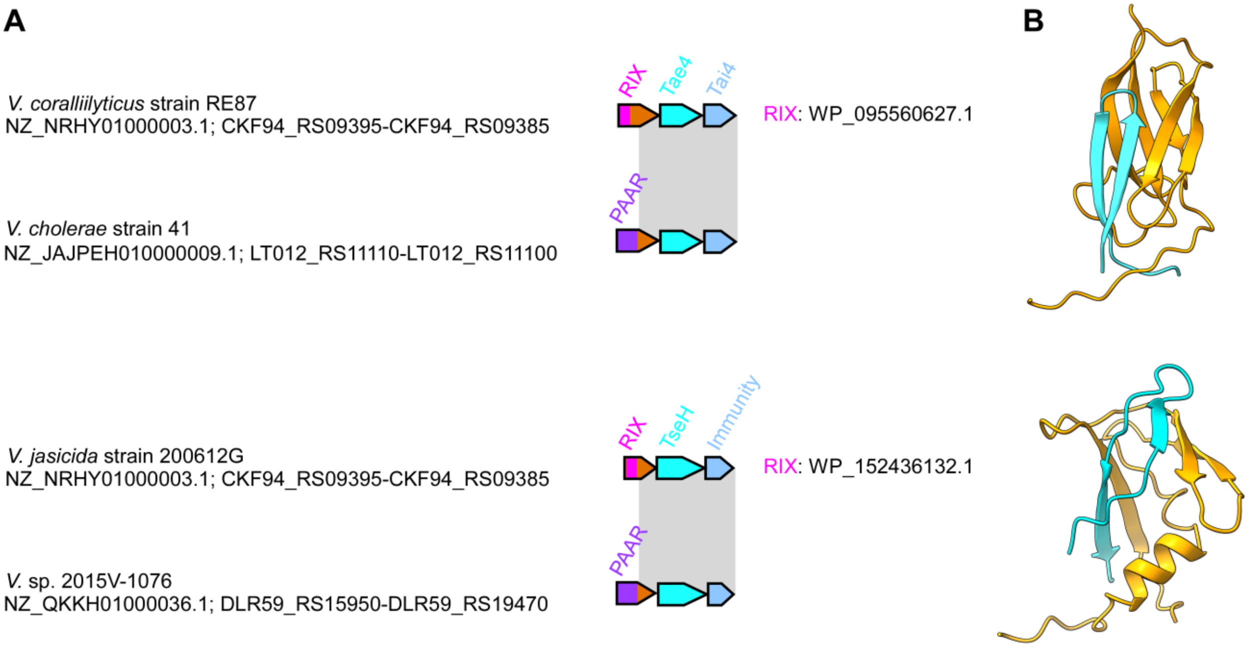


**Supplementary Fig. S8. RIX-containing proteins can serve as T6SS tethers. (A)** The gene structure of the operons encoding WP_095560627.1 in *V. coralliilyticus* RE87 and WP_152436132.1 in *V. jasicida* 200612G. Operons encoding a module homologous to the C-terminal extension of the RIX-containing protein and the downstream-encoded effector and immunity pair are shown below; a gray rectangle denotes the region of homology. The strain names, the GenBank accession numbers, and the locus tags are provided. Genes are denoted by arrows indicating the direction of transcription. The names of encoded proteins or domains are denoted above. **(B)** Interactions between the RIX-containing proteins and their downstream-encoded effector were predicted by AlphaFold2. The interaction interfaces, corresponding to amino acids 102-185 of WP_095560627.1 (orange) and 1-21 of Tae4 (cyan) (top), and amino acids 76-134 of WP_152436132.1 (orange) and 1-24 of TseH (cyan) (bottom), are shown.

### Supplementary Datasets

**Supplementary Dataset S1.** RIX domain-containing proteins identified in this study.

**Supplementary Dataset S2.** The presence of T6SS in RIX domain-encoding bacterial genomes.

### Supplementary Files

**Supplementary File S1.** AlphaFold2-generated PDB files used in this study.

### Supplementary Tables

**Supplementary Table S1. A list of bacterial strains used in this study.**

| Strain name | Genotype | Comments | Source |
|---|---|---|---|
| *Vibrio parahaemolyticus* BB22OP | Wild type | Used for generating deletion strains | Obtained from Kim Orth; [1] |
| *Vibrio parahaemolyticus* BB22OP Δ*hns*/Δ*tme1* | Δ*vpbb_rs05425*/ Δ*vpbb_rs15030* | BB22OP derivative containing in-frame deletions of *hns* and *tme1*. Used for secretion assays | [2] |
| *Vibrio parahaemolyticus* BB22OP Δ*hns*/Δ*tme1*/Δ*hcp1* | Δ*vpbb_rs05425*/ Δ*vpbb_rs15030*/ Δ*vpbb_rs06665* | BB22OP derivative containing in-frame deletions of *hns, tme1,* and *hcp1*. Used for secretion assays | [2] |
| *Pseudomonas aeruginosa* PAO1 | Wild type | Used as a template to amplify the *tse1* CDS | Obtained from Avigdor Eldar |
| *Vibrio parahaemolyticus* RIMD 2210633 | Wild type | Used for generating deletion strains | Obtained from Kim Orth |
| *Vibrio parahaemolyticus* RIMD 2210633 Δ*hns*/Δ*tdhAS*/ Δ*vp_rs06130*/ Δ*vp_rs06745*/ *vp_rs06875*$^{AAA}$<br><br>(Surrogate attacker) | Δ*vp_rs05510*/Δ*tdhAS*/ Δ*vp_rs06130*/ Δ*vp_rs06745*/ *vp_rs06875*$^{AAA}$ | RIMD 2210633 derivative with a constitutively active, effectorless T6SS1. Used as a surrogate platform in competition assays and in secretion assays | [3] |
| *Vibrio parahaemolyticus* RIMD 2210633 Δ*hns*/Δ*tdhAS*/ Δ*vp_rs06130*/ Δ*vp_rs06745*/ *vp_rs06875*$^{AAA}$/Δ*hcp1* | Surrogate attacker Δ*hcp1* | Surrogate attacker derivative with an inactive T6SS1. Used as a surrogate platform in competition | [3] |

| | | | |
|---|---|---|---|
| | | assays and in secretion assays | |
| *Vibrio natriegens* ATCC 14048 | Wild type | Used as prey in competition assays | ATCC collection |
| *Vibrio coralliilyticus* ATCC BAA-450 | Wild type | Used for generating deletion strains, as the attacker in competition assays, and in secretion assays | ATCC collection |
| *Vibrio coralliilyticus* ATCC BAA-450 Δ*hcp1* | Δ*vic_rs16330* | ATCC BAA-450 derivative containing an in-frame deletion of *vic_rs16330*; used as the attacker in competition assays, and in secretion assays | [3] |
| *Vibrio campbellii* ATCC 25920 | Wild type | Used for cloning | ATCC collection |
| *Escherichia coli* DH5α (λ pir) | K-12 derivative laboratory strain containing λ pir | Used for plasmid maintenance and cloning | Obtained from Eric V. Stabb |
| *Escherichia coli* BL21 (DE3) | Laboratory strain | Used for protein expression and toxicity assays | Obtained from Kim Orth |

**Supplementary Table S2. A list of plasmids used in this study.**

| Plasmid name | Description | Purpose | Source |
|---|---|---|---|
| pBAD$^K$/Myc-His | pBR322 ori-containing plasmid harboring a Kan$^R$ cassette, *araC*, and an MCS following a P*bad* promoter | Used for arabinose-inducible expression | [4] |
| pRti1$^M$ | pBAD$^K$/Myc-His plasmid containing the CDS of WP_050778602.1 from *V. coralliilyticus* ATCC BAA-450 in frame with a C-terminal Myc-His tag | Used for the arabinose-inducible expression of WP_050778602.1 in *E. coli* and *V. parahaemolyticus* | This study |

| | | | |
|---|---|---|---|
| pPER5 | pBAD$^{K}$/Myc-His with a PelB signal peptide inserted at the 5' end of the MCS | Used for the arabinose-inducible expression of proteins targeted to the periplasm in *E. coli* | [5] |
| pTme1$^{peri}$ | pPER5 plasmid containing the CDS of Tme1 (WP_015297525.1) from *V. parahaemolyticus* BB22OP in frame with an N-terminal PelB signal peptide and the C-terminal Myc-His tag | Used for the arabinose-inducible expression of Tme1 protein targeted to the periplasm in *E. coli* | [2] |
| pTme1$^{61\text{-}310/peri}$ | pPER5 plasmid containing the CDS of Tme1$^{61\text{-}310}$ (truncation of the first 60 amino acids of Tme1) in frame with an N-terminal PelB signal peptide and a C-terminal Myc-His tag | Used for the arabinose-inducible expression of Tme1$^{61\text{-}310}$ protein targeted to the periplasm in *E. coli* | This study |
| pTme1$^{1\text{-}225/peri}$ | pPER5 plasmid containing the CDS of Tme1$^{1\text{-}225}$ (truncation of amino acids 226-310 of Tme1) in frame with an N-terminal PelB signal peptide and a C-terminal Myc-His tag | Used for the arabinose-inducible expression of Tme1$^{1\text{-}225}$ protein targeted to the periplasm in *E. coli* | This study |
| pBAD33.1$^{F}$ | pBAD33.1 with a FLAG tag inserted at the 3' end of the MCS | Used for the arabinose-inducible expression of proteins | [2] |
| pTme1 | pBAD33.1$^{F}$ plasmid containing CDS of Tme1 in frame with the C-terminal FLAG tag of the plasmid | Used for the arabinose-inducible expression of proteins in *V. parahaemolyticus* BB22OP | [2] |
| pTme1$^{61\text{-}310}$ | pBAD33.1$^{F}$ plasmid containing CDS of Tme1$^{61\text{-}310}$ (truncation of the | Used for the arabinose-inducible expression of proteins | This study |

| | | | |
|---|---|---|---|
| | first 60 amino acids of Tme1) in frame with the C-terminal FLAG tag of the plasmid | in *V. parahaemolyticus* BB22OP | |
| pTme1$^{1-111}$-Tse1 | pBAD33.1$^{F}$ plasmid containing CDS for Tme1$^{1-111}$ (the first 111 amino acids of Tme1) fused to Tse1 (WP_003088027.1) from *Pseudomonas aeruginosa* PAO1, in frame with the C-terminal FLAG tag of the plasmid | Used for the arabinose-inducible expression of proteins in *V. parahaemolyticus* BB22OP | This study |
| pTse1 | pBAD33.1$^{F}$ plasmid containing CDS of Tse1 in frame with the C-terminal FLAG tag of the plasmid | Used for the arabinose-inducible expression of proteins in *V. parahaemolyticus* BB22OP | This study |
| pWP_005536620.1 | pBAD33.1$^{F}$ plasmid containing CDS of WP_005536620.1 from *V. campbellii* ATCC 25920 in frame with the C-terminal FLAG tag of the plasmid | Used for the arabinose-inducible expression of proteins in *E. coli* and *V. parahaemolyticus* | This study |
| pWP_005530005.1 | pBAD33.1$^{F}$ plasmid containing CDS of WP_005530005.1 from *V. campbellii* ATCC 25920 in frame with the C-terminal FLAG tag of the plasmid | Used for the arabinose-inducible expression of proteins in *E. coli* and *V. parahaemolyticus* | This study |
| pWP_038863399.1 | pBAD33.1$^{F}$ plasmid containing CDS of WP_038863399.1 from *V. campbellii* ATCC 25920 in frame with the C-terminal FLAG tag of the plasmid | Used for the arabinose-inducible expression of proteins in *E. coli* and *V. parahaemolyticus* | This study |
| pEffector | pBAD33.1$^{F}$ plasmid containing CDS of WP_005534959.1 | Used for the arabinose-inducible expression of proteins | This study |

| | | | |
|---|---|---|---|
| | from *V. campbellii* ATCC 25920 in frame with the C-terminal FLAG tag of the plasmid | in *E. coli* and *V. parahaemolyticus* | |
| pEffector +Immunity | pBAD33.1$^{F}$ plasmid containing CDS of WP_005534959.1 and WP_005534960.1 from *V. campbellii* ATCC 25920 in frame with the C-terminal FLAG tag of the plasmid | Used for the arabinose-inducible expression of proteins in *E. coli* and *V. parahaemolyticus* | This study |
| pImmunity | pBAD33.1$^{F}$ plasmid containing CDS of WP_005534960.1 in frame with the C-terminal FLAG tag of the plasmid | Used for the arabinose-inducible expression of proteins in *E. coli* and *V. parahaemolyticus* | This study |
| pRIX | pBAD33.1$^{F}$ plasmid containing CDS of WP_157622110.1 from *V. coralliilyticus* ATCC BAA-450 in frame with the C-terminal FLAG tag of the plasmid | Used for the arabinose-inducible expression of proteins in *V. coralliilyticus* and *V. parahaemolyticus* | This study |
| pRte1 | pBAD33.1$^{F}$ plasmid containing CDS of WP_006958655.1 from *V. coralliilyticus* ATCC BAA-450 in frame with the C-terminal FLAG tag of the plasmid | Used for the arabinose-inducible expression of proteins in *E. coli* | This study |
| pRte1$^{H94A}$ | pBAD33.1$^{F}$ plasmid containing CDS of WP_006958655.1 from *V. coralliilyticus* ATCC BAA-450 with a substitution of histidine 94 for alanine, in frame with the C-terminal | Used for the arabinose-inducible expression of proteins in *E. coli* and the secretion assay in *V. parahaemolyticus* | This study |

| | | | |
|---|---|---|---|
| | FLAG tag of the plasmid | | |
| pRte1$^{S106A}$ | pBAD33.1$^{F}$ plasmid containing CDS of WP_006958655.1 from *V. coralliilyticus* ATCC BAA-450 with a substitution of serine 106 with alanine, in frame with the C-terminal FLAG tag of the plasmid | Used for the arabinose-inducible expression of proteins in *E. coli* | This study |
| pRIX+Rtei1 | pBAD33.1$^{F}$ plasmid containing CDS of WP_157622110.1, WP_006958655.1, and WP_050778602.1 from *V. coralliilyticus* ATCC BAA-450 in frame with the C-terminal FLAG tag of the plasmid | Used for the arabinose-inducible expression of proteins in *V. parahaemolyticus* POR1 | This study |
| pRIX+Rte1$^{H94A}$ | pBAD33.1$^{F}$ plasmid containing CDS of WP_157622110.1 and WP_006958655.1 with a H94A mutation from *V. coralliilyticus* ATCC BAA-450 in frame with the C-terminal FLAG tag of the plasmid | Used for the arabinose-inducible expression of proteins in *V. parahaemolyticus* POR1 | This study |
| pRtei1 | pBAD33.1$^{F}$ plasmid containing CDS of WP_006958655.1 and WP_050778602.1 from *V. coralliilyticus* ATCC BAA-450 in frame with the C-terminal FLAG tag of the plasmid | Used for the arabinose-inducible expression of proteins in *V. parahaemolyticus* POR1 | This study |

| | | | |
|---|---|---|---|
| pGML10 | pGML10 *E. coli-S. cerevisiae* shuttle vector, GAL1-10 promoter with a Myc tag at the 3' end of the MCS | Used for the galactose-inducible expression of proteins in *S. cerevisiae* | Riken |
| pGML10:eGFP | pGML10 containing the CDS of enhanced GFP in the EcoRI site of the MCS, in frame with the C-terminal Myc tag of the plasmid | Used for the galactose-inducible expression of eGFP in *S. cerevisiae* | This study |
| pGML10:WP_005536620.1 | pGML10 containing the CDS of WP_005536620.1 in frame with the C-terminal Myc tag of the plasmid | Used for the galactose-inducible expression of proteins in *S. cerevisiae* | This study |
| pGML10:WP_005530005.1 | pGML10 containing the CDS of WP_005530005.1 in frame with the C-terminal Myc tag of the plasmid | Used for the galactose-inducible expression of proteins in *S. cerevisiae* | This study |
| pGML10:WP_038863399.1 | pGML10 containing the CDS of WP_038863399.1 in frame with the C-terminal Myc tag of the plasmid | Used for the galactose-inducible expression of proteins in *S. cerevisiae* | This study |
